# Building gene regulatory networks from scATAC-seq and scRNA-seq using Linked Self-Organizing Maps

**DOI:** 10.1101/438937

**Authors:** Camden Jansen, Ricardo N. Ramirez, Nicole C. El-Ali, David Gomez-Cabrero, Jesper Tegner, Matthias Merkenschlager, Ana Conesa, Ali Mortazavi

## Abstract

Rapid advances in single-cell assays have outpaced methods for analysis of those data types. Different single-cell assays show extensive variation in sensitivity and signal to noise levels. In particular, scATAC-seq generates extremely sparse and noisy datasets. Existing methods developed to analyze this data require cells amenable to pseudo-time analysis or require datasets with drastically different cell-types. We describe a novel approach using self-organizing maps (SOM) to link scATAC-seq and scRNA-seq data that overcomes these challenges and can generate draft regulatory networks. Our SOMatic package generates chromatin and gene expression SOMs separately and combines them using a linking function. We applied SOMatic on a mouse pre-B cell differentiation time-course using controlled Ikaros over-expression to recover gene ontology enrichments, identify motifs in genomic regions showing similar single-cell profiles, and generate a gene regulatory network that both recovers known interactions and predicts new Ikaros targets during the differentiation process. The ability of linked SOMs to detect emergent properties from multiple types of highly-dimensional genomic data with very different signal properties opens new avenues for integrative analysis of single-cells.

## Introduction

The ability to analyze hundreds to thousands of individual cells using new functional sequencing assays has revolutionized the current state of scientific and biomedical research^1^. For example, single-cell gene expression studies have allowed the identification of rare cell populations in a variety of samples ranging from immune cell systems^2^ to circulating tumor cells^3^. Comprehensive atlases of gene expression are being built for tissues such as the Drosophila brain throughout its lifespan^4^ to an entire mouse^5^. Inspired by the wealth of new insights from single-cell RNA-seq, there has been a plethora of single cell genomic technologies developed in the last few year (reviewed in^6^). For example, single-cell profiling of chromatin accessibility^7–9^ has generated a lot of excitement because of the wealth of insights generated within large scale surveys of chromatin accessibility and gene regulation through projects like ENCODE^10^.

However, unlike single-cell RNA-seq, chromatin accessibility mapping from individual cells yields sparse information of the open chromatin landscape^11, 12^ due to the intrinsic limitation of numbers of chromosomes per nucleus. It has been difficult for previous analysis platforms to handle the scarcity and noise inherent in data of this type.

Recently, a number of tools have been developed to try and combat this issue. chromVAR^13^ uses cells with the highest proportion of reads to build a model of the expected number of fragments per total reads for every respective motif site in the genome, and computes deviation scores from this model to cluster single-cells. This method, while effective, requires the generation of a list of transcription factor binding sites through mass motif scanning which, in this work, necessitated the loosening of strict Type I error control and the creation of a custom, well-curated list of transcription factor motifs. Another application, scABC^13^, manages to cluster cells of different cell-types well by using the total cell accessibility signal to provide weights to an unsupervised clustering of the cells using K-mediods and thus identifies landmark regions that are only open in each found population. The cells are then re-clustered using the respective landmarks. However, this technique would likely become confused by time course data from the same cell-type as it may be too similar to generate proper landmarks.

More recent techniques attempt to correct for the scarcity of scATAC-seq data by leveraging imputed pseudo-time orderings^14^. For example, Cicero^15^ uses the ordering of cells to make small aggregate pools before computing correlations. Alternatively in a study of human hematopoietic cell differentiation, Buenrostro and colleagues^16^ also assigned pseudotime ordering so that accessibility peaks could be smoothed by a lowess function. Both of these methods make extensive use of pseudotime orderings, and thus, require systems that have a strong differentiation lineage (with preferably known markers). Here we introduce a method for jointly analyzing scRNA-seq and scATAC-seq data that cannot be ordered by pseudotime by taking a “gene/region-centric” approach using self-organizing maps.

Self-organizing maps (SOMs) are a type of artificial neural networks, also referred to as a Kohonen network^17, 18^(Supp. Fig. 1). SOMs are trained using unsupervised learning to generate a low-dimensional representation of data and can be visualized using two-dimensional maps, which allows for a low-dimensional representation of this high-dimensional data. Individual SOM nodes (or neurons) have a weight vector that is in the same dimension as the input data vectors and neighboring nodes on a SOM reflect similarities across the input data space vector. Thus, trained SOMs provide an intuitive platform for identifying clusters in high-dimensional datasets. For example, SOMs trained on gene expression data or chromatin data^19^ from multiple cell types in human and mouse have identified complex relationships across high-dimensional genomic data^10, 20–22^. Additionally, SOMs have been used to structure and interrogate the transcriptome in single-cells during cellular reprogramming^23^. SOMs provide a natural visual and powerful platform for the analysis and integration of high-dimensional data of different types.

**Figure 1.**
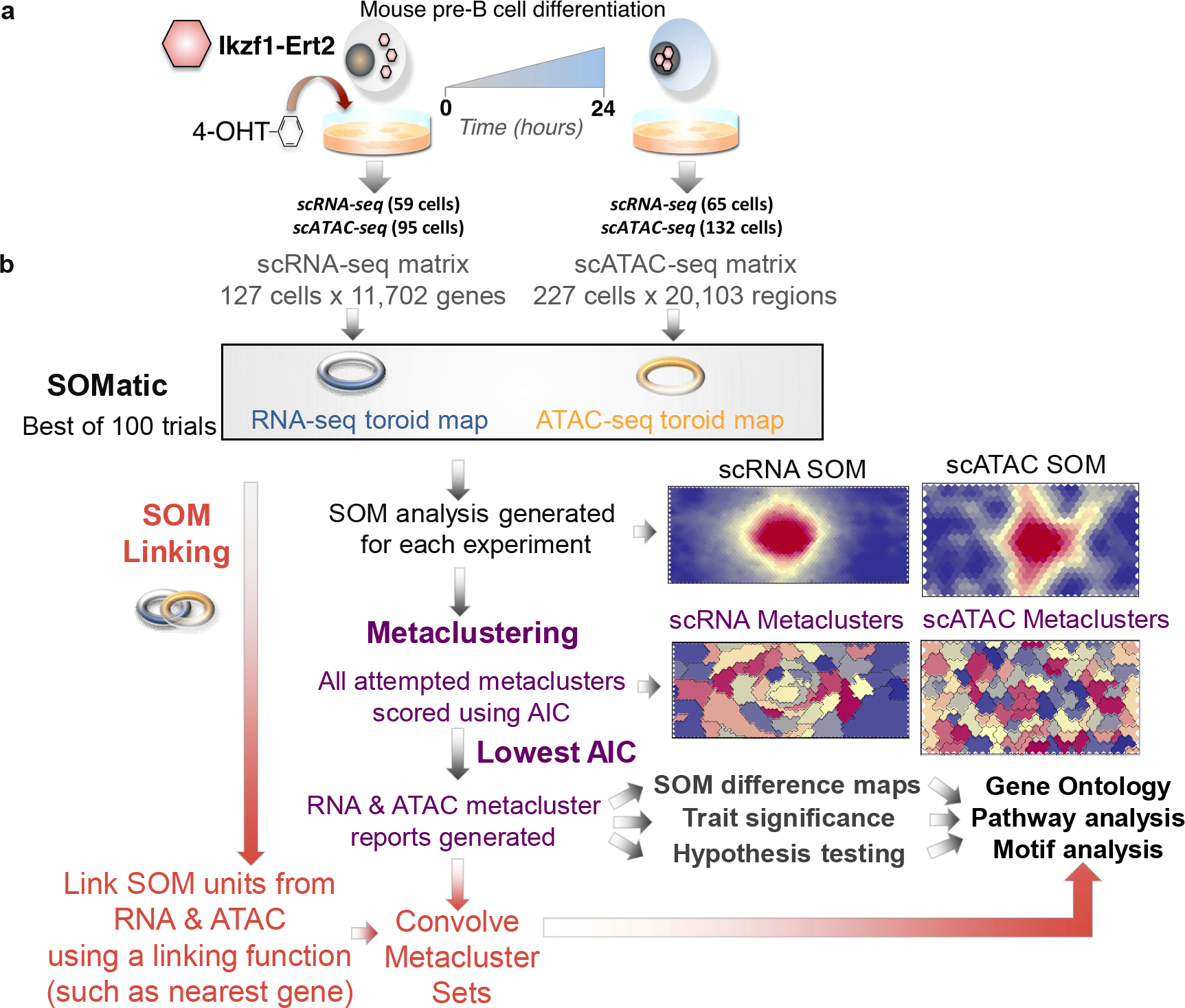
Single-cell multi-data integration using SOMs. (a) An inducible Ikzf1 mouse pre-B cell-line was used to track changes in gene expression and chromatin accessibility during differentiation (0 and 24-hours) in single-cells. (b) Single-cell RNA-seq and ATAC-seq data from an inducible mouse pre-B cell-line were independently trained using SOMatic to generate single-cell SOMs and metaclustered using AIC scoring. These clusters were convolved with the new SOM fusion algorithm to generate pair-wise metaclusters of chromatin regions with similar profiles across the single-cell dataset that regulate genes that also share similar profiles. These pair-wise clusters were mined for regulatory connections through motif enrichment analysis.

As part of our work in the STATegra consortium (STATegra.eu), we performed single-cell RNA-seq and single-cell ATAC-seq using a mouse pre-B cell model system^24^ during cellular differentiation. This system provides a high-resolution view into a narrow transition in pre-B cell development, whereby we induce cell differentiation in response to a sudden doubling of Ikaros expression. Our data only contains two time points and represents a fairly drastic change in chromatin accessibility and gene expression over that period, and thus, would be a poor candidate for pseudo-time analysis. In addition, this data is sufficiently sparse and noisy to give even powerful algorithms like UMAP^25^ difficulty from a gene or genome region perspective (Supp. Fig. 2, 3).

**Figure 2.**
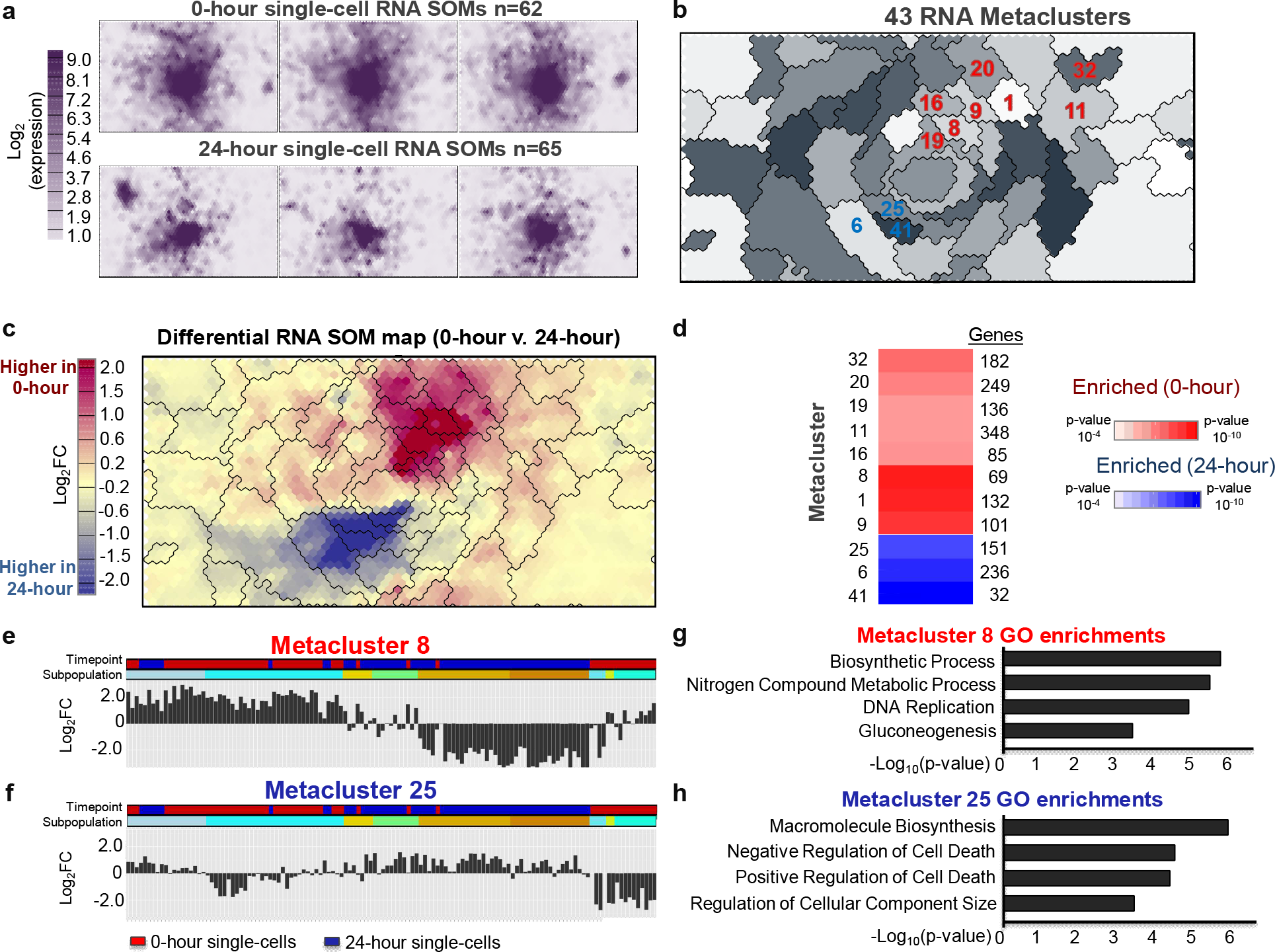
Single-cell gene expression patterns during cellular differentiation are profiled using SOMatic. (a) A SOM was generated for the single-cell RNA-seq dataset (0-hour 62 cells, 24-hour 65 cells). Maps for 3 cells from each time point were arbitrarily selected for display. (b) 43 metaclusters were identified using AIC scoring. Metacluster number and color were arbitrarily assigned for visualization purposes. (c) SOM difference map comparing 0-hour and 24-hour time-points. Maps for cells from 0 and 24-hour timepoints were averaged to generate a single map for each and then subtracted to create a map that represented gene expression fold change during pre-B cell development. Overlaid metacluster divisions generally follow contours of the map. (d) Trait enrichment analysis deployed on gene metaclusters revealed which are enriched in each time point. Metaclusters of interest are highlighted in panel b. (e-f) Summary showing the representative expression profile for metaclusters 8 and 25. Columns are individual cells color-coded for 0 and 24-hour time-points ordered by hierarchical clustering on every metacluster representative gene expression profile. Cell subpopulations are represented by a 40% cut on that clustering. (f-g) Top gene ontology terms for the 69 genes in metacluster 8 and the 151 genes in metacluster 25.

**Figure 3.**
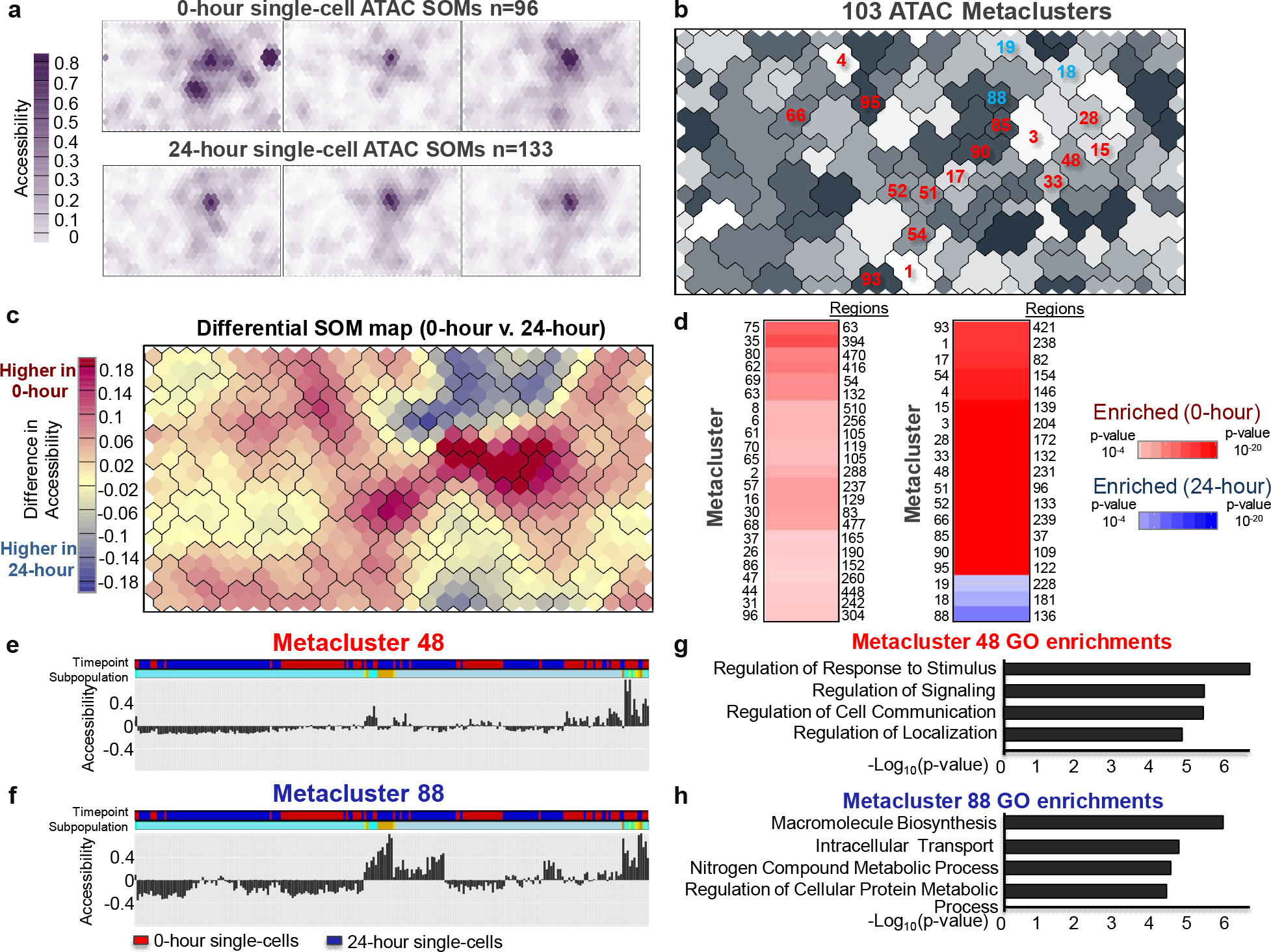
SOMatic reveals the dynamic chromatin landscape in single-cells. (a) A chromatin SOM was generated for the single-cell ATAC-seq dataset (0-hour 96 cells, 24-hour 133 cells). Maps for 3 cells from each timepoint were arbitrarily selected for display. (b) 103 metaclusters were identified using AIC scoring. Metacluster number and color were arbitrarily assigned for visualization purposes. (c) SOM difference map comparing 0-hour and 24-hour time-points. Maps for cells from 0 and 24-hour timepoints were averaged to generate a single map for each and then subtracted to create a map that represented chromatin accessibility fold change during pre-B cell development. Overlaid metacluster divisions generally follow contours of the map. (d) Trait enrichment analysis deployed on gene metaclusters revealed which are enriched in each time point. Metaclusters of interest are highlighted in panel b. (e-f) Summary showing the representative accessibility profile for SOM metaclusters 48 and 88. Columns are individual cells color-coded for 0 and 24-hour time-points ordered by hierarchical clustering on every metacluster representative gene expression profile. Cell subpopulations are represented by a 40% cut on that clustering. (f-e) Top gene ontology terms for genes associated to chromatin elements from SOM metaclusters 48 and 88. Association was determined through use of the GREAT algorithm (See methods).

We used SOMatic to create two SOMs in order to identify significant groups of expressed genes and chromatin elements that jointly change during the time course. The two SOMs were then linked using a novel algorithm to find metaclusters of genes and associated genomic regions that show similar profiles during pre-B cell differentiation. The regulatory regions in these clusters were mined for enriched motifs that allowed us to infer a predicted regulatory network downstream of Ikaros. Our flexible and comprehensive approach is first of its kind to provide an analysis platform that combines these different single-cell data types without leveraging cell ordering and effectively identifies regulatory programs.

## Results

### Integration of single-cell data types using SOM

In order to study changes in gene expression and chromatin accessibility for single-cells, we utilized an inducible pre-B model system^24^ and performed single-cell RNA-seq and single-cell ATAC-seq before and after cellular differentiation (Experimental methods). The goal was to link the data from these methods in a meaningful way to study individual genome region/gene interactions, and this was accomplished by developing the computational pipeline shown in Figure 1. We began by training separate self-organizing maps (SOMs) for each dataset. The result is a set of SOM units that contain genes and genome regions that have a very similar signal profile across each of the single cells at both time points (Summary maps in Supp. Fig. 4). To reduce the signal dropout and technical noise prevalent in single cell data, our SOM analysis tool produces clusters of these units, called metaclusters^19^, which maintain the SOM’s scaffold topology by only combining adjacent units and contain similar gene expression and chromatin accessibility profiles. Finally we combine the patterns found in each SOM using a pipeline that links metaclusters from both gene expression and chromatin accessibility. These linked metaclusters (LM) contain sets of chromatin regions that have similar open chromatin signal profiles that are in the proximity of genes that also share a similar profile (although not necessarily the same profile in RNA and ATAC) and can be mined using gene ontology, pathway analysis, and motif discovery. Our method easily extends a traditional single data-type analysis to one that focuses on the integration of fundamentally different data like single-cell RNA-seq and ATAC-seq in order to recover evidence of co-regulation.

**Figure 4.**
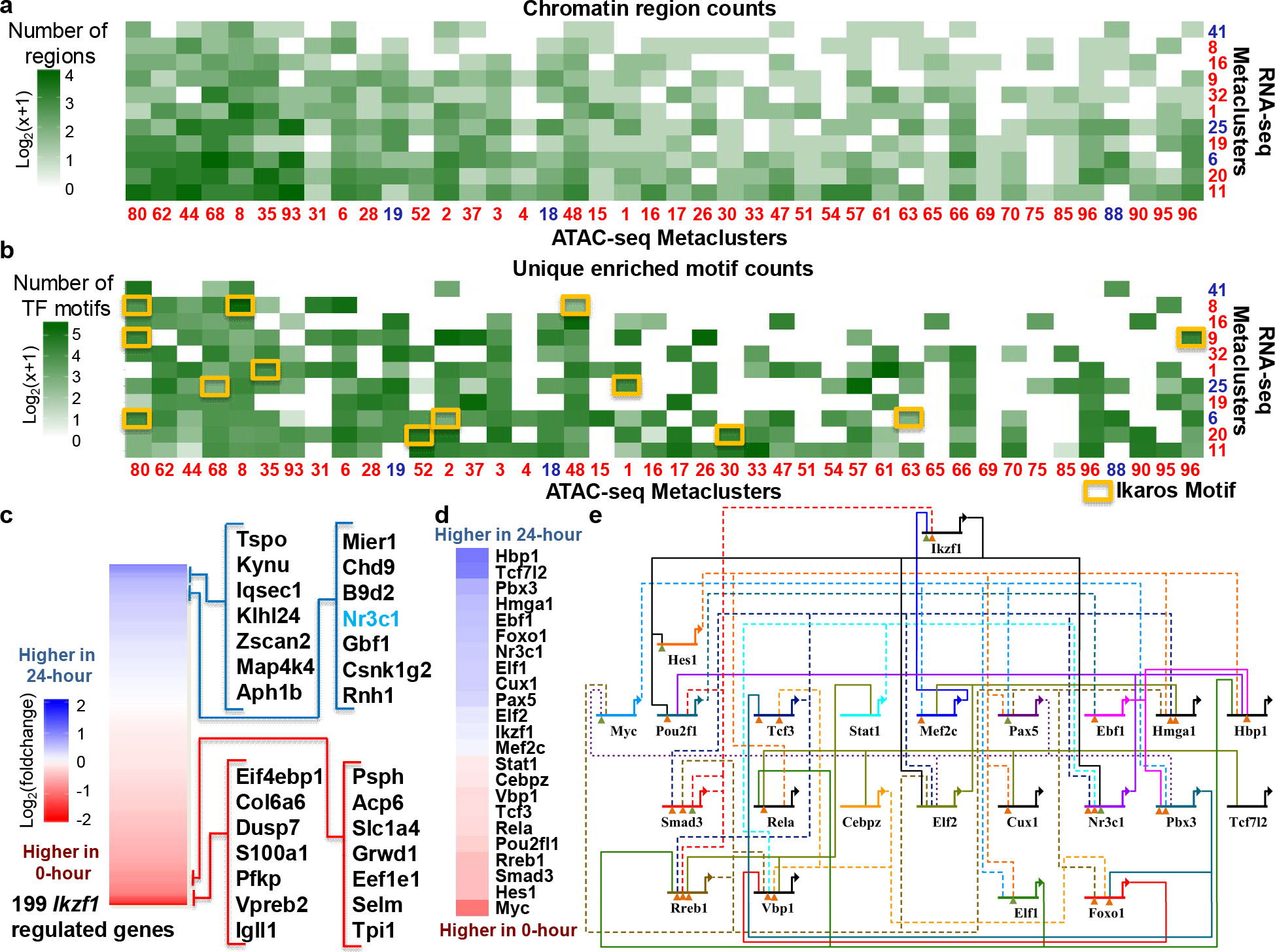
Transcriptional regulation by *Ikzf1* recovered using SOM fusion. (a) Size of pair-wise metaclusters that contain both differentially-expressed genes and differentially-accessible chromatin sites. Metaclusters of genes and regions with a higher enrichment at 24-hours are colored blue and are ordered to place larger pair-wise metaclusters in the bottom left. (b) Number of statistically-significant motifs found in each pair-wise metacluster from (a). Presence of the *Ikzf1* motif in the pair-wise metacluster is noted. (c) Heatmap of expression fold change for genes predicted to be regulated by *Ikzf1*. Genes with the largest change between time points are noted. Nr3c1 is the only listed gene that is labeled as a transcription factor. (d) Predicted downstream targets of *Ikzf1* with significant change over the time course. Each gene is labels with the fold change between time points with the same scale as 4c. (e) Predicted gene regulatory network downstream of *Ikzf1*. Genes are ordered left to right by their fold change over the time course. Connections are dashed if their signal is significantly lower at the 24-hour time point. Connections at each gene are labeled by level of evidence found in existing literature. Green triangles indicate experimental evidence and orange triangles indicate previous computational prediction.

### Identification of dynamic gene expression metaclusters

We trained a 40 × 60 SOM on the 127 scRNA-seq datasets (62 single-cells for 0-hour; 65 single-cells for 24-hour) using 11,702 genes that had expression greater than 1 FPKM in at least 5% of cells. As expected, slices of this map (Fig. 2a), which correspond to single cells, show a general reduction of gene expression over time. SOMatic identified 43 RNA metaclusters that reflect the various gene expression profiles present in the data (Fig. 2b). We validated that these metaclusters were properly determined by calculating the UMatrix and density map for this SOM and overlaying the metacluster boundaries on top of these maps (Supp. Fig. 5) for visual inspection. The metaclusters followed the breaks in these maps as expected and thus provide a robust representation of the different profiles present in the single-cell data.

One of the strengths of the SOM approach is that we can perform logical operations on the feature maps. We computed a map by averaging maps from the cells in each time point and subtracting them to determine which metaclusters reflect meaningful gene expression differences across time (Fig. 2c). We performed a correlation analysis to determine which metaclusters were consistently enriched across the cells in each time-point as previously described^19^. We found statistically-significant differences across time in 11 RNA metaclusters, 8 of which were enriched in 0-hour and 3 in 24-hour (Fig. 2d, p-value <10^−4^-10^−10^). For example, RNA metacluster 8 consists of 21 SOM units and contains 69 genes enriched in 0-hour single-cells such as *Igll1* and *Vpreb1* (Fig. 2e). Similarly, metacluster 25 consists of 33 units and contains 151 genes enriched in 24-hour cells such as *Mier1* and *Foxp1*, which has been shown to control mature B-cell survival in mice^26^ (Fig. 2e). Gene ontology analysis revealed a series of genes enriched for negative regulation of apoptosis in 24-hour cells, while DNA replication genes were represented in 0-hour cells (Fig. 2f). This is consistent with the transition of gene programs necessary for coordinating pre-B cell differentiation^27^.

### Mapping the pre-B single-cell chromatin landscape architecture using SOMs

We performed single-cell ATAC-seq^8^ with a total of 229 cells passing our quality controls to explore the change in chromatin accessibility over the differentiation time-course. We recovered on average 53,864 unique chromatin fragments per cell (Supp. Fig. 6e). Using peaks taken from a set of pooled ATAC-seq experiments over three biological replicates with 50,000 cells for each time-point, we quantified the ATAC-seq signal in these peaks for each cell. We built a data matrix from chromatin regions detected in at least 2% of cells (5 cells) for a total 20,103 ATAC-seq peaks due to the sparse nature of single-cell ATAC-seq.

A 20 × 30 SOM was trained on this scATAC data matrix. Similar to the RNA SOM, scATAC feature maps (Fig. 3a) revealed a general closing of the chromatin in 24-hour cells, which is normal for cells undergoing differentiation. Clustering the units from this SOM resulted in the identification of 103 chromatin metaclusters (Fig. 3b). Visual inspection of these clusters confirmed that these clusters properly follow the breaks in the UMatrix and density map (Supp. Fig. 7).

A SOM difference map and hypothesis analysis for all 103 chromatin metaclusters revealed 39 metaclusters that exhibit open chromatin signal in 0-hour cells and 3 metaclusters in with higher signal in the 24-hour cells (Fig. 3c-d). Gene ontology enrichments for genes in the vicinity of the regions from two of the most significant metaclusters (Fig 3e), 48 (0-hour enriched; 231 peaks) and 88 (24-hour enriched; 136 peaks), reveal that these genes are enriched for cell signaling and DNA replication programs as predicted (Fig. 3f). Thus, SOMs are capable of revealing patterns of chromatin accessibility from sparse single-cell ATAC-seq data in a dynamic model system.

### Application of multi-omic single-cell data integration using Linked SOMs

Cellular differentiation occurs as a consequence of dynamics in expression of networks of genes controlled by cis-regulatory elements, which must be open in order to function properly. The linker pipeline within SOMatic attempts to convolve the metaclusters from RNA and chromatin accessibility SOMs in order to interrogate the dynamics of the system. In brief, the pipeline subsets chromatin regions within the same chromatin metacluster into linked metaclusters (LM) using the expression of the gene whose regulatory region (using the same algorithm as GREAT^28^) overlaps the element. Thus, if a set of regions are in a LM, these regions share a similar chromatin accessibility profile and are in the vicinity of genes that also share a similar gene expression profile (See Supp. Fig. 8 for an overview). This coherence of joint profiles gives a much higher expectation that these regions will be similarly regulated than grouping on accessibility or gene expression alone.

We applied this new pipeline to our scRNA and scATAC SOMs and analyzed a total of 103 × 43 = 4,429 LMs to identify 462 LMs that were significantly dynamic in both chromatin accessibility and their nearby genes (Fig 4a). Based on our assumption that these LMs were similarly regulated, we mined each LM separately for known transcription factor binding site motifs using FIMO with a q-value cutoff of .05. This generated ~4.1 million candidate motifs, which is substantially more than results from motif analysis on bulk data:less than 50k and 500k for peaks and enriched peaks respectively (Supp. Fig. 9); random LMs also gave us fewer candidate motifs, with an average of ~1.46 million motif positions in 100 trials (Supp. Fig. 10). Additionally, to determine enrichment, LMs with a percentage of regions containing each transcription factor motif that was significantly (pvalue < .05) enriched over the baseline were reported, (Fig. 4b), reducing the ~4.1 million candidate motifs to 112,550 high-confidence potential gene regulatory network connections or 3,480 high-confidence active transcription factor/active transcription factor connections.

The differentiation of this B3 cell line is initiated by a doubling the amount of Ikaros in the nucleus of each cell and we therefore focused our analysis on Ikaros as the root node of a gene regulatory network. Several of the LMs are enriched for the Ikaros motif, including 12 of the LMs showing both RNA expression and chromatin accessibility differences. In total, we found 199 genes, with 205 nearby potential cis-regulatory regions that contain the motif, that may be regulated directly by Ikaros (Fig. 4c), including genes known to be differentially expressed in this system, such as *Igll1* (Supp. Fig. 11) and *Vpreb2*^29^ as well as the transcription factor *Nr3c1*^30^. This factor has been previously implicated as being downstream of Ikaros and was the only transcription factor (with a motif) in the list of the top 30 differentially-expressed genes. To validate these connections, *Ikzf1* ChIP data^27^ was interrogated at the same 0hr and 24hr time points at each of the 205 potential cis-regulatory regions. Of these, 200 (~98%) of these regions had *Ikzf1* ChIP signal (more than 1 RPKM) in one or both of the time points and 79 (~39%) had a significant change over the time course (more than 2x fold change). Loci for the 4 transcription factors predicted to be regulated by Ikaros were further visually inspected and each of the nearby potential cis-regulatory regions had a significant change over the time course (Supp. Fig. 12-15).

We built a gene regulatory network of transcription factors that we predicted were connected to Ikaros to identify indirect, secondary changes to gene expression as a direct result of changes in Ikaros concentration at the direct targets TFs. This network is tied directly to the model system in that it only uses genome segments that are open in either time-point. We determined which factors downstream of Ikaros showed a significant change in expression across the time-series (Fig. 4d) and determined the connections between them (Fig. 4e). Each of these genes has been shown to be important in B-cell differentiation. For example, the activation of *Hbp1* has been shown to prevent c-Myc-mediated transcription^31^ and, together with a down-regulation of *Myc* expression, stops B-cell proliferation. The temporal enrichment of predicted targets downstream of *Myc* can be found in Supp. Fig. 16.

About 15% of connections in this network have been previously described^30, 32–35^, which include *Mef2c* to *Ikaros*^36^ and *Pax5* and *Myc*’s negative feedback loop^37, 38^, or have been previously computationally predicted^39, 40^(50%), and we identify new connections like *Rreb1* to *Myc*(~35%)(Supp. Fig. 17). The identification of both direct and indirect regulation from a sudden doubling of Ikaros demonstrates the power of the Linked SOMs for analyzing highly-dimensional multi-omics data.

## Discussion

In this work, we used a gene- and chromatin-centric analysis using SOMs on a mouse pre-B time-course data of single-cell RNA-seq and ATAC-seq separately and, then, convolved them to find synergistic effects. Combining the metaclusters from multiple SOMs as a pair-wise set generates a data-space that combines the properties from both without any assumptions about how the data relates to each-other. Due to the inheritance of each SOM’s properties, the linked metaclusters (LMs) contain genome regions that should be similarly regulated: not only is the chromatin accessibility of those regions similar across the cells, but the nearby genes they regulate share expression patterns. Thus, these LMs can be mined for motif enrichment and return a higher number of significant motif sites than simply dividing the data set randomly or by signal changes in either data set separately.

We used this SOM linking technique to explore the regulatory control of the lymphoid regulator *Ikzf1* during one step of B-cell development. 12 LMs enriched in the *Ikzf1* motif contained regions that had similarly-differential chromatin accessibility between time points and had had differentially expressed genes. Our analysis successfully recovers known biology about *Ikzf1* regulation on target genes *Igll1*, *Vpreb2*, and *Nr3c1* and novel regulatory information through discovery of possible downstream mechanisms for B-cell activation. Following the interactions around the network provides many exciting, new avenues for research.

It is important to note, however, that these predicted regulatory connections use an extremely stringent statistical cutoff to be as confident as possible, and thus, do not recover some of the linkages predicted based on *Ikzf1* ChIP data^27^ such as Ikaros’s involvement in the regulation of *Myc* and *Foxo1*. While we do detect these connections at an early portion of the pipeline, the genome sequence in those regulatory regions are too different from the canonical motif to pass our stringent filters. *Foxo1* had an Ikaros motif in an open chromatin region near its transcription start site, but the motif only had a q-value of 0.112, which was far above the threshold.

Our approach for combining multi-omic data through linked SOMs is amenable to integrating other single-cell technologies for the purpose of multi-omic data analysis as long as a linking function can be found. For example, the profiling of small RNAs, such as miRNAs^41^, in single cells could be linked with a standard scRNA-seq experiment through the use of target prediction algorithms. The hypothetical LMs in that case would include groups of miRNAs with similar expression patterns such that their target RNA also has similar expression patterns. Following identification of these groups, functional analysis could be done on each group target RNAs and these functions could be passed back to the miRNA in the group. This is just one example of an exciting experimental and computational design that linked SOMs enable. The ability to perform multi-omic experiments from a single-cell is now achievable for several biochemical and genomic platforms^42–45^ with more being developed every day. We foresee the ability to connect the patterns in multi-omic data using algorithms like linked SOMs to be integral in using this new technology to the fullest.

## Methods

### Pre-B cell differentiation

ERt2-Ikaros inducible B3 cells were cultured in Iscove’s Modified Dulbecco’s Medium (IMDM) supplemented with 10% FBS. Differentiation was induced as previously shown^27^. Briefly, cells were induced with 20mM of 4-hydroxytamoxifen (4OHT), over the course of 24 hours. Prior to performing single-cell experiments, cells were washed twice with cold 1X PBS.

### Single-cell RNA-seq

Single cells were isolated using the Fluidigm C1 System. Single cell C1 runs were completed using the smallest IFC (5-10 um) based on the estimated size of B3 cells. Briefly, cells were collected for 0 and 24-hour time-points at a concentration of 400 cells/μl in a total of 50 μl. To optimize cell capture rates on the C1, buoyancy estimates were optimized prior to each run. Each individual C1 capture site was visually inspected to ensure single-cell capture and cell viability. After visualization, the IFC was loaded with Clontech SMARTer kit lysis, RT, and PCR amplification reagents. After harvesting, cDNA was normalized across all libraries from 0.1-0.3 ng/μl and libraries were constructed using Illumina’s Nextera XT library prep kit per Fluidigm’s protocol. Constructed libraries were multiplexed and purified using AMPure beads. The final multiplexed single-cell library was analyzed on an Agilent 2100 Bioanalyzer for fragment distribution and quantified using Kapa Biosystem’s universal library quantification kit. The library was normalized to 2 nM and sequenced as 75bp paired-end dual indexed reads using Illumina’s NextSeq 500 system at a depth of ~1.0-2.0 million reads per library.

### Single-cell ATAC-seq

Single-cell ATAC-seq was performed using the Fluidigm C1 system as done previously^8^. Briefly, cells were collected for 0 and 24-hours post treatment with tamoxifen, at a concentration of 500 cells/μl in a total of 30-50 μl. Additionally, 3 biological replicates of ~50,000 cells were collected for each measured time-point to generate bulk ATAC-seq measurements. Bulk ATAC-seq was performed as previously described^46^. ATAC-seq peak calling was performed using bulk ATAC-seq samples. ATAC-seq peaks were then used to estimate single-cell ATAC-seq signal. Our C1 single-cell capture efficiency was ~70-80% for our pre-B system. Each individual C1 capture site was visually inspected to ensure single-cell capture. In brief, amplified transposed DNA was collected from all captured single-cells and dual-indexing library preparation was performed. After PCR amplification of single-cell libraries, all subsequent libraries were pooled and purified using a single MinElute PCR purification (Qiagen). The pooled library was run on a Bioanalyzer and normalized using Kappa library quantification kit prior to sequencing. A single pooled library was sequenced as 40bp paired-end dual indexed reads using the high-output (75 cycle) kit on the NextSeq 500 from Illumina. Two C1 runs were performed for 0 and 24-hour single-cell ATAC-seq experiments.

### Single-cell RNA-seq data processing

Single-cell RNA-seq libraries were mapped with Tophat^47^ to the mouse Ensembl gene annotations and mm10 reference genome. Single-cell libraries with a mapping rate less than 50% and less than 450,000 mapped reads were excluded from any downstream analysis. Analysis was performed using 0 and 24-hour single-cells. Cufflinks^48^ version 2.2.1 was used to quantify expression from single-cell libraries using Cuffquant. Gene expression measurements for each single-cell library were merged and normalized into a single data matrix using Cuffnorm.

### Bulk and single-cell ATAC-seq data processing

Single-cell libraries were mapped with Bowtie^49^ to the mm10 reference genome using the following parameters (bowtie -S -p 2 --trim3 10 −X 2000). Duplicate fragments were removed using Picard (http://picard.sourceforge.net) as previously performed^8^. We considered single-cell libraries that recovered > 5k fragments after mapping and duplication removal. Bulk ATAC-seq replicates were mapped to the mm10 reference genome using the following parameters (bowtie -S --trim3 10 -p 32 -m 3 -k 1 -v 2 --best -X 2000). Peak calling was performed on bulk replicates using HOMER with the following parameters (findPeaks <tags> -o <output> -localSize 50000 -size 150 -minDist 50 -fragLength 0). The intersection of peaks in three biological replicates was performed. A consolidated list of peaks was generated from the union of peaks from 0 and 24 hour time-points.

### ChIP-seq analysis

*Ikzf1* ChIP-seq data for 0 and 24-hour pre-B cells^24^ was mapped to the mm10 reference genome using Bowtie2^50^. For all samples, we filtered duplicated reads and those with a mapping quality score below 20. To identify peaks, we used the CLCbio Peak Finder software<u>_ENREF_38</u>^51^ with default parameters and control input libraries. We defined significant peaks with an adjusted p-value <0.01 also using biological replicates.

### Training and Metaclustering of the individual RNA and ATAC SOMs

We use the SOMatic package, which is a combination of tools written in C++ and R designed for the analysis and visualization of multidimensional genomic or gene expression data, to train our individual SOMs. The SOMatic package also builds a customized, optional javascript viewer to mine the results visually. Installation information for this package can be found at https://github.com/csjansen/SOMatic.

For the RNA-seq SOM, we built a matrix of 11,702 expressed genes in 127 single cells and we used half the genes (5851) to train a self-organizing map with a toroid topology with size 40 × 60 with 5,851,000 million time steps (1000 epochs) as previously described^20^ to select the best of 100 trials based on lowest fitting error. The entire matrix was used for scoring this best trial to generate the final SOM. The SOMatic website for this SOM can be viewed at http://crick.bio.uci.edu/STATegra/RNASOM/

Similarly, the ATAC-seq data was organized into a matrix consisting of scATAC signal in 227 cells at 20,103 ATAC-seq peaks (from pooled data) and half of the peaks were used to train a SOM with a toroid topology with size 40 × 60 using 19,955,000 time steps (1000 epochs) as previously described^20^. The best of 100 trials based on lowest fitting error was selected and the entire matrix was used for scoring the final SOM. The SOMatic website for this SOM can be viewed at http://crick.bio.uci.edu/STATegra/ATACSOM/

SOM units with similar profiles across cells were grouped into metaclusters^19, 20^ using SOMatic. Briefly, metaclustering was performed using k-means clustering to determine centroids for groups of units. Metaclusters were built around these centroids so that each cluster is in one piece to maintain the SOM topology. SOMatic’s metaclustering function attempts all metacluster numbers within a range given and scores them based on Akaike information criterion (AIC)^52^. The penalty term for this score is calculated using a parameter called the “dimensionality,” which is the number of independent dimensions in the data. We performed a hierarchical clustering on the SOM unti vectors and counted the number of clusters that were present at a height level equal to 30% of the total distance in the clustering. For the ATAC-seq SOM, the dimensionality was calculated to be 37, and for the RNA-seq SOM, the dimensionality was calculated to be 62.

For the RNA metaclustering, we tried all k between 20 and 50, whereas for the ATAC metaclustering we tried all k between 80 and 120. The metacluster number with the lowest AIC score was the one chosen for each SOM. For ATAC-seq, 103 metaclusters had the best score, and for RNA-seq, 43 metaclusters had the best score. R scripts for generating metacluster reports are provided in the SOMatic package. Metatcluster/Trait correlation and hypothesis testing analysis were done as previously described^19^.

### Hyperparameter Variation

There are inherent trade-offs that have to be kept in mind when choosing SOM parameters for training and metaclustering. For example, the size of a SOM is typically one of the most important decisions to be made in analyses of this type. A smaller SOM may group elements together that do not belong together and will reduce the statistical power of down-stream analysis, and a larger SOM may separate elements that belong in the same cluster but are separated due to noise, causing down-stream analysis to miss patterns that may exist. Similarly, the number of timesteps and the learning rate will change the chances of under and over-clustering by changing how the SOM scaffold morphs into the topology of the data. Proper metaclustering can improve the robustness of the SOM by easily revealing improper training due to poor parameters.

The scRNA-seq SOM was built with additional sizes 20 × 30 and 80 × 120 with little change to the calculated number of metaclusters, with 41 and 44 respectively. The 20 × 30 SOM was not chosen for the final analysis due to the occurrence of multiple 1-unit metaclusters, which indicates an underclustering. The 80 × 120 SOM was not chosen due to having a metacluster that contained a unit in each row which indicated a possible overclustering. The number of timesteps and learning rate chosen were determined to be sufficient due to the smoothness of the final summary map (Supp. Fig. 4a). An insufficient value in either of these parameters would cause the summary to have large breaks in total signal between neighboring units, indicating under-training.

The scATAC-seq SOM was also built with sizes 10×15 and 40×60 with little change to the calculated number of metaclusters, with 95 and 117 respectively. Again, the smaller SOM was not chosen for the final analysis due to multiple 1-unit metaclusters, which indicates an underclustering. The 40 × 60 SOM was not chosen due to the map focusing too much on regions that were unique to each cell, indicating overclustering. The number of timesteps and learning rate chosen were determined to be sufficient due to the smoothness of the final summary map (Supp. Fig. 4b).

### Linked SOMs

In order to define this, a few preliminary definitions are required. For a set *A* of data vectors, it is possible to define a set of *n* vectors, *B*, indexed on a 2D lattice to partition *A* into *n* subsets with each vector assigned to the subset *i* iff *B*_*i*_ is the closest element of *B* to that vector. Due to the 2D indexing lattice that they are placed on, each vector in *B* is adjacent to its closest member in *B*, with “closest” defined by an unsupervised neural network. The set of vectors, *B*, is the set of SOM units.

Similarly, it is possible to define a set of *m* vectors, *M*, to partition *B* into *m* subsets, *S*, with each vector assigned to the subset *i* iff *M*_*i*_ is the closest element in *M* to that vector such that a path can be drawn on the lattice using only elements of *S_i_*. This path requirement is in place to maintain the SOM topology calculated in training of the neural network. The subsets, *S*, are the metaclusters defined previously.

Let *G* be the set of gene vectors from a number of RNA-seq experiments and let *R* be the set of genome region vectors defined by ATAC-seq peaks. Using the procedure above, it is possible to segment these sets into metaclusters, named *N* and *M* respectively. Between these two metacluster partitions, we can define a linker mapping, *h*, from *R* to *G*. Using a linker mapping designed to link the individual SOM datatypes, we can define a set of partitions, *F^M,N,h^*, where (*r,g*) ϵ (*R,G*) is an element of *F^M,N,h^_ij_* iff *h*(*r*)=*g, g* ϵ *N_j_*, and *r* ϵ *M*_*i*_. In this case, the linker mapping that we use to link RNA and open chromatin data is an implementation of the GREAT^28^ OneClosest algorithm with a cutoff of 50kb to build regulatory regions around transcription start sites for each gene and check if these regions overlap with the ATAC-seq peaks. The resulting Linked SOM metaclusters (LMs) contain clusters of similar genome regions such that their linked genes are also similar.

### Motif Analysis

The regulatory regions in each Linked SOM metacluster were separately scanned for motifs from the HOCOMOCOv11 mouse motif database^53^ with FIMO v4.9.0_4^54^ using a q-value threshold of .05. Then, for each transcription factor in the database, the percentage of regions in each LM with a motif for that factor was calculated. To determine enrichment, the percentages for each transcript factor were separately compared in a one-tailed z-score analysis. LMs with a percentage that was significantly (pvalue < .05) enriched over the baseline, the average percentage across all LMs for that transcription factor motif, was reported for each transcription factor. Finally, transcription factors with a statistically significant number of motifs were mapped to the gene fused to the regulatory region the motif was found within. The full list of these potential connections can be found here: http://crick.bio.uci.edu/STATegra/LinkedMotifMappings.txt.

### GEO Accession

GEO accession number for data is GSE89285.

---

**Supplementary Figure 1.**
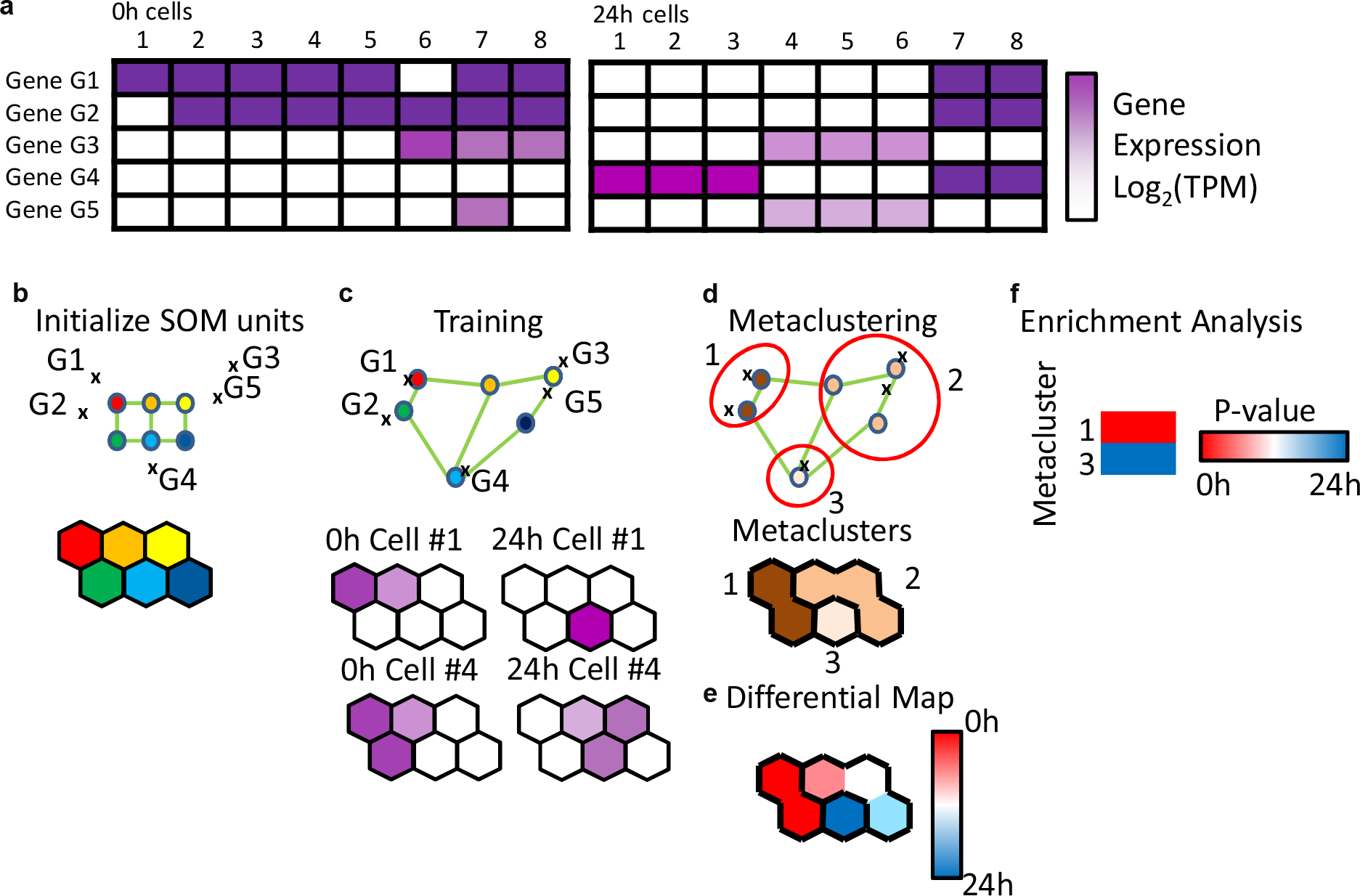
Self-Organizing Map Clustering Overview. (a) Example heatmap for 5 genes’ expression in a typical single-cell RNA-seq with 2 time points. Genes G1 and G2 are enriched at 0h with two 0h cells missing that signal due to technical noise and gene G4 is enriched at 24hr. Genes G3 and G5 also have a similar expression pattern with two cells missing signal in G5 due to technical noise, but are not particularly enriched in either time point. (b) 2D representation of the genes’ expression profile with an initial SOM scaffold. The colors in the scaffold correspond to those the map below. (c) 2D representation of the genes’ expression profile with a typical trained SOM scaffold overlaid. The maps below represent the signal for each unit in the labeled experiment’s dimension. For example, only gene G4 has signal in 24h Cell #1, and thus, only the unit near G4 has signal on the map. (d) Neighboring units with similar expression profiles are metaclustered to fix the overclustering of genes G1 and G2 into separate units. (e) Multiple individual maps can be combined into one through arithmetic. This map represents the average of each 24h map subtracted from the average of each 0h map. (f) Trait enrichment analysis can be applied on each metacluster to provide a p-value for enrichment in a particular time point. Here, metacluster 1, containing genes G1 and G2, is enriched in 0h, and metacluster 3, containing gene G3, is enriched in 24h.

**Supplementary Figure 2.**
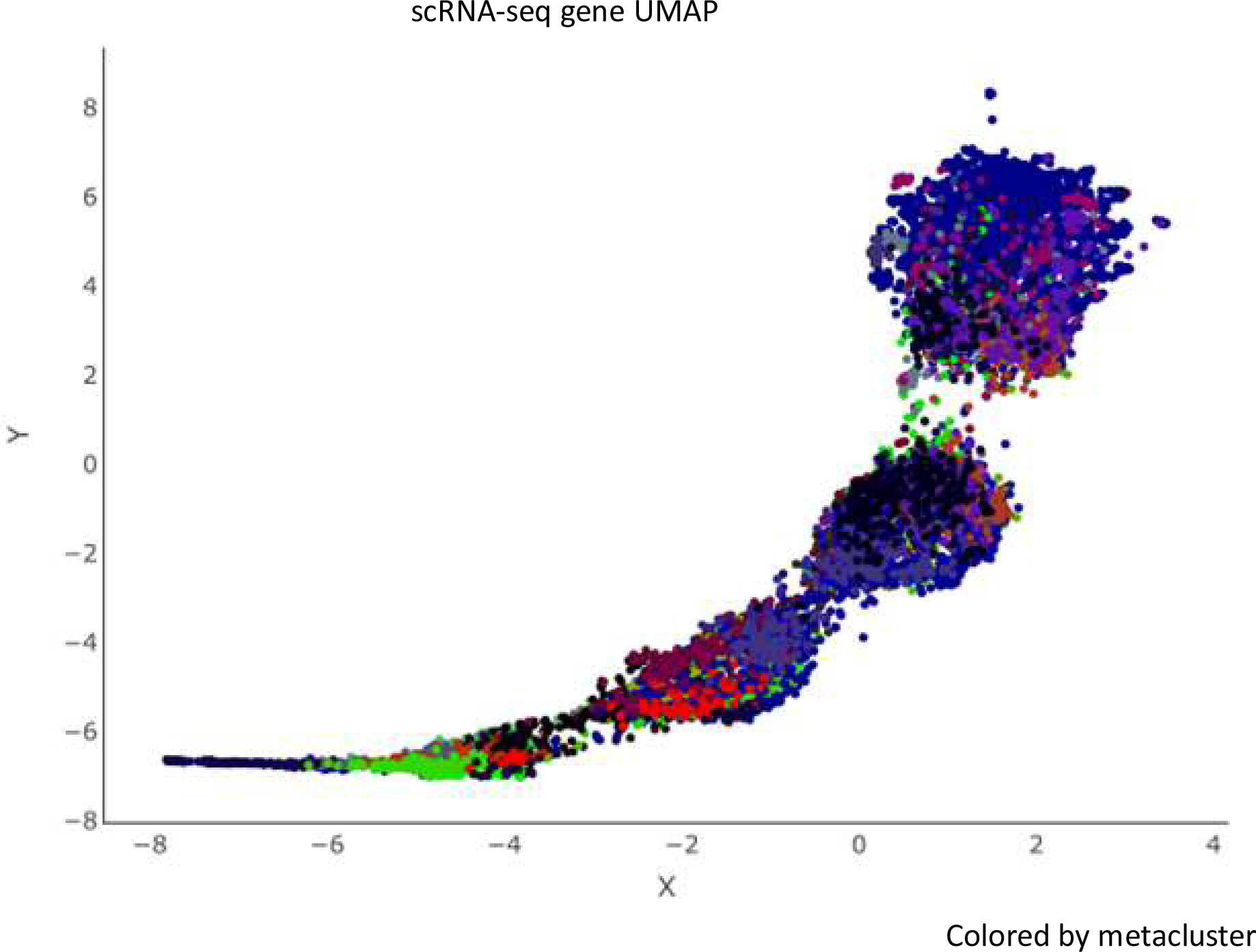
scRNA-seq gene UMAP. UMAP^25^ generated using uwat^55^ from scRNA-seq data with each point representing a gene’s expression in each cell. The umap is separated into 4 large clusters, which provides a poor level of resolution for downstream analysis. Points were colored by RNA SOM metacluster, which divides the large clusters into many sub-clusters.

**Supplementary Figure 3.**
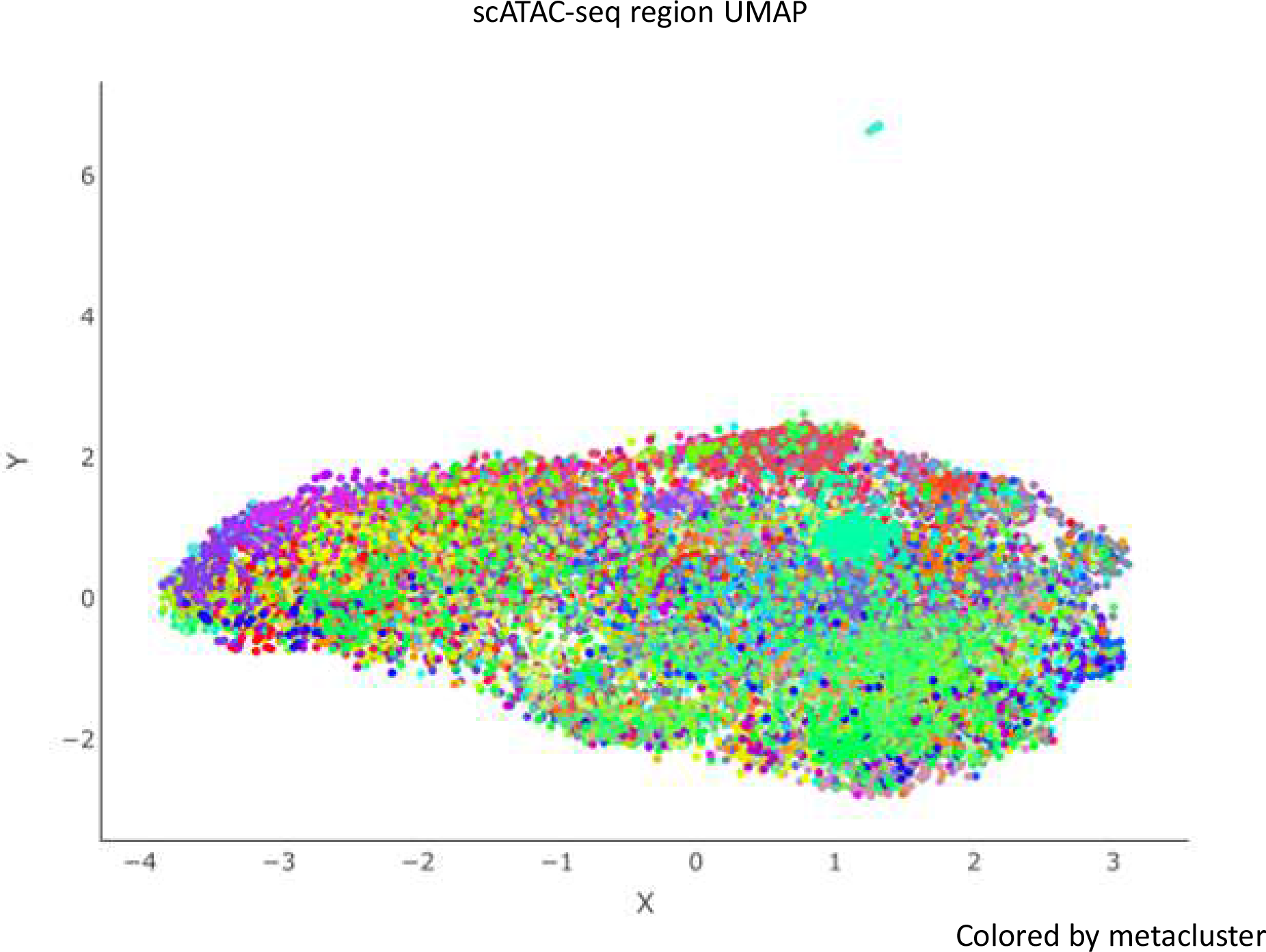
scATAC-seq region UMAP. UMAP^25^ generated using uwat^55^ from scATAC-seq data with each point representing a genome region’s ATAC-seq signal in each cell. The umap could not be separated into any significant clusters. Points were colored by ATAC SOM metacluster, which divides the large cluster into many sub-clusters.

**Supplementary Figure 4.**
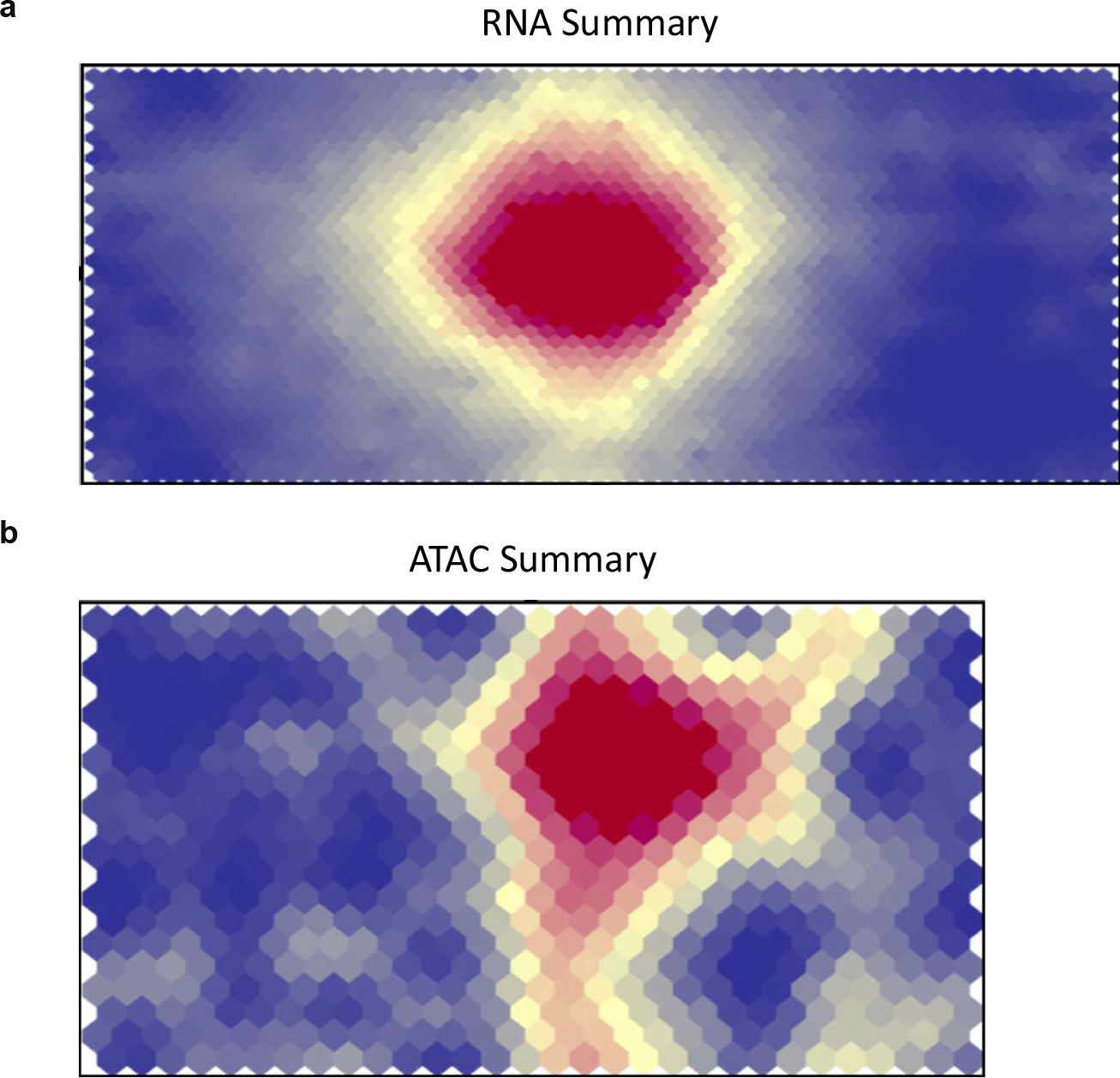
SOM summary maps (total signal in every cell) (a-b) Summary maps for the (a) RNA and (b) ATAC SOMs. Each unit’s value is generated by totaling the values in the full SOM unit’s vector. A blue-white-red color spectrum was used. These graphs are mainly used to determine ‘smoothness’ of the SOM fit and to see if more timesteps or changes to the learning rate are needed.

**Supplementary Figure 5.**
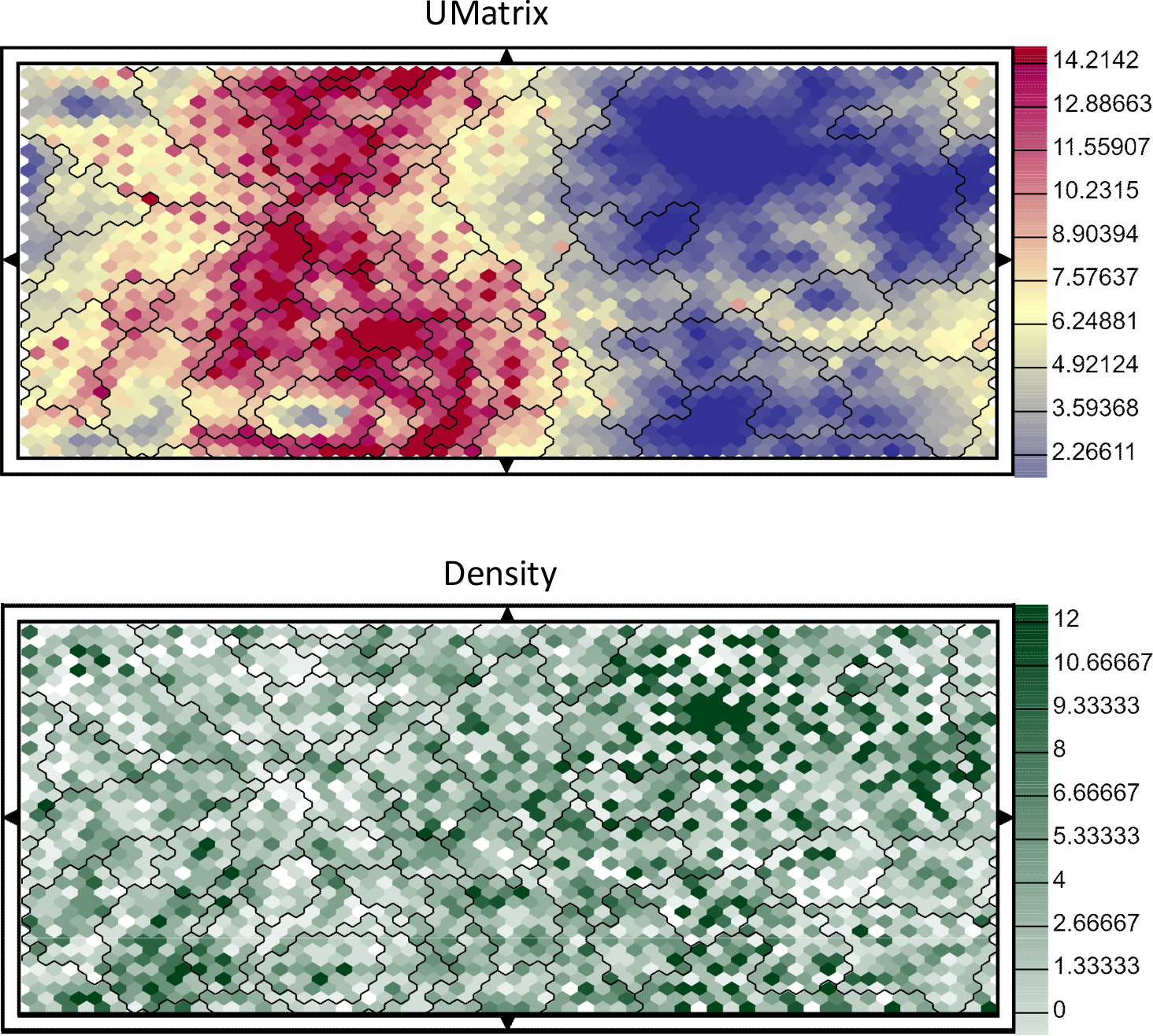
Statistic maps for scRNA-seq SOM. (a) U-Matrix for the SOM built with the single-cell RNA-seq dataset. Each unit contains the average of the distance to all neighboring units. Metacluster divisions are overlaid. Areas of high distance correspond primarily to a metacluster division. (b) Density map for the RNA-seq SOM. The color corresponds to the number of genes found in each unit. Metacluster divisions are overlaid. Most metaclusters are ruled by a few high density units.

**Supplementary Figure 6.**
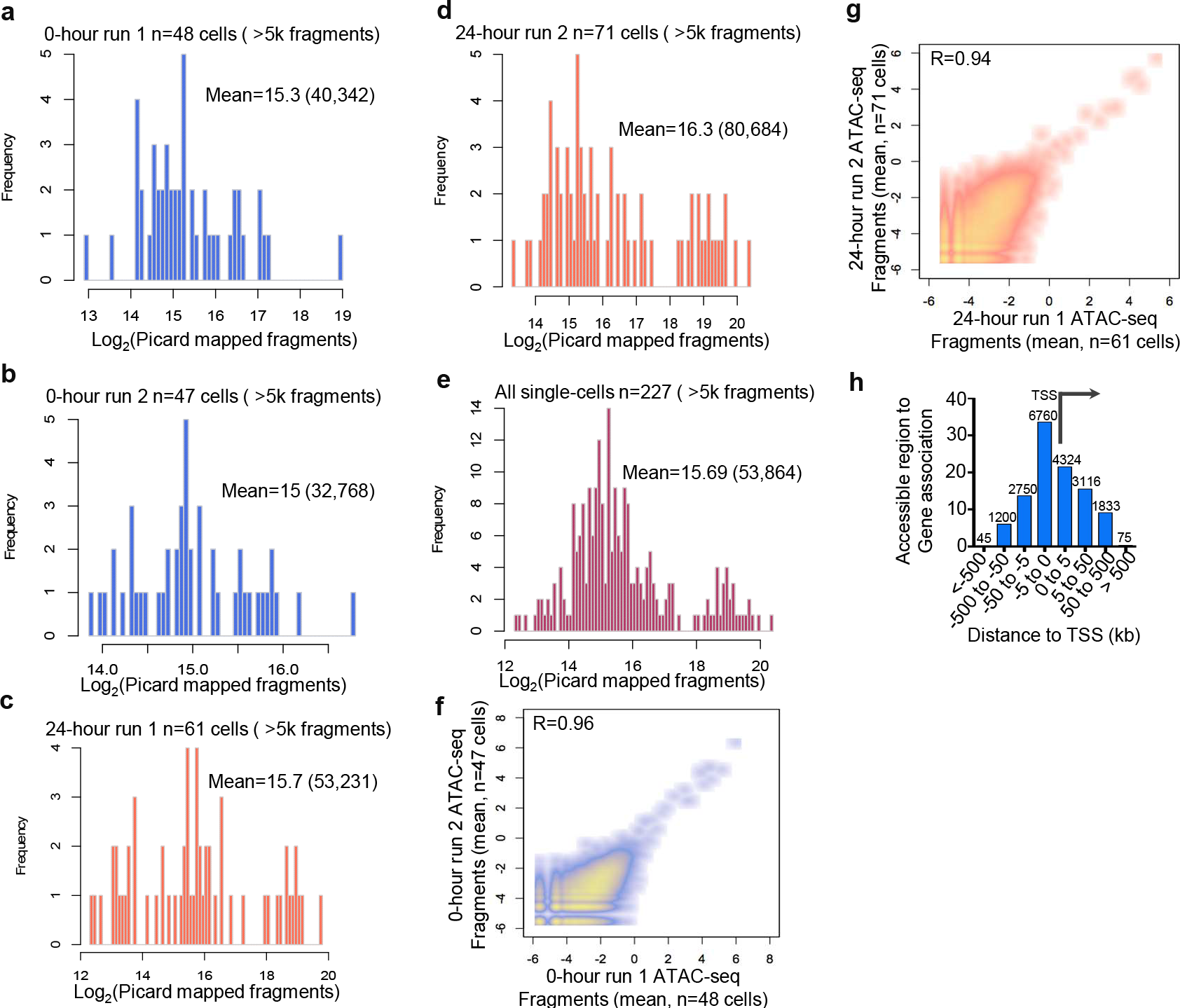
Single-cell ATAC-seq library statistics. (a-b) 0-hour ATAC-seq Picard mapped fragment distribution for single-cell libraries between two C1 runs. (c-d) 24-hour ATAC-seq Picard mapped fragment distribution for single-cell libraries between two C1 runs. (e) ATAC-seq Picard mapped fragment distribution of all 227 single-cell libraries. (f-g) Pearson correlation between C1 runs for 0 and 24-hour data sets. Averaged ATAC-seq fragments were determined for each C1 run. (h) Distribution of 20,103 ATAC-seq regions to nearest single gene TSS.

**Supplementary Figure 7.**
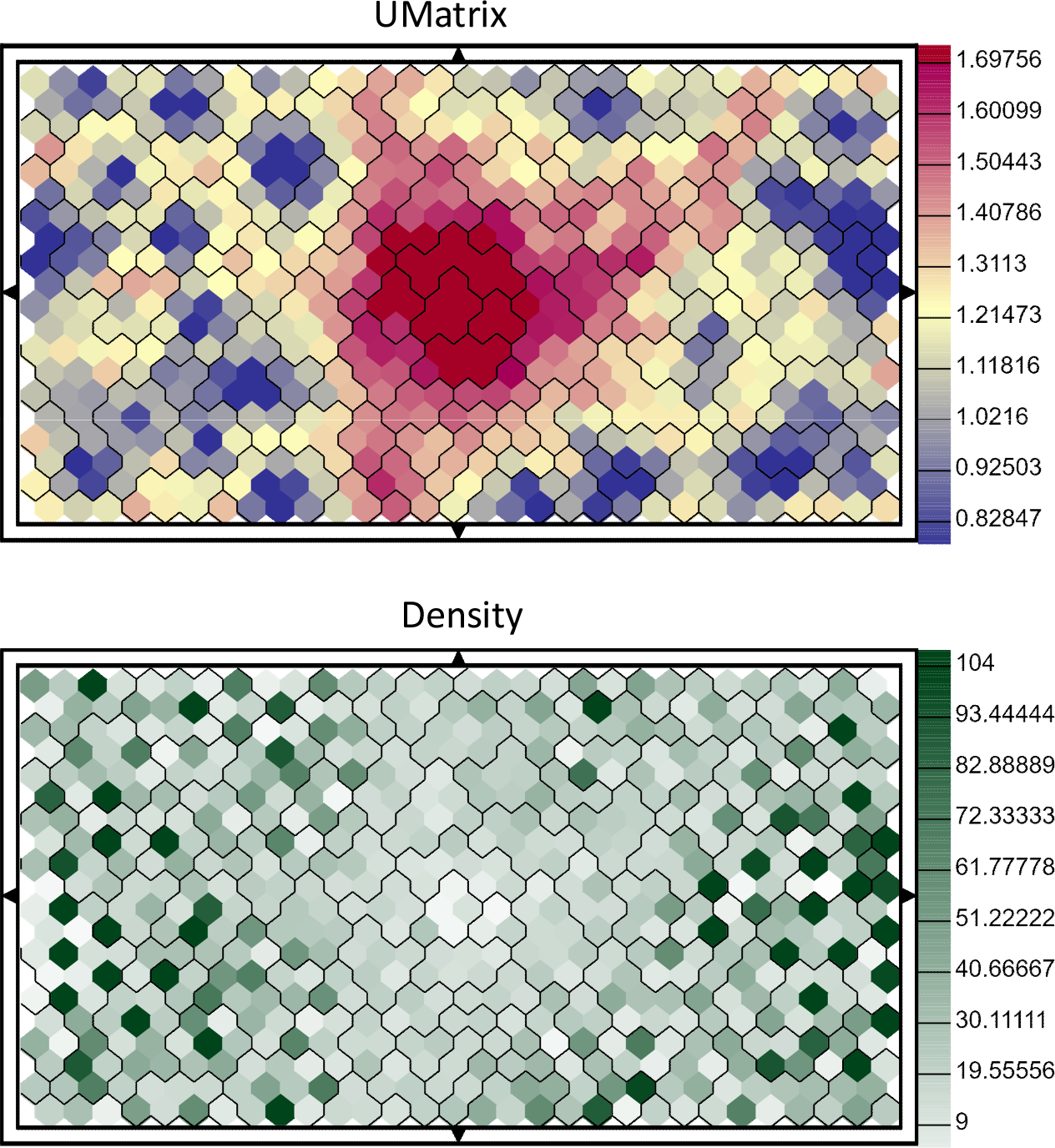
Statistic maps for scATAC-seq SOM. (a) U-Matrix for the SOM built with the single-cell ATAC-seq dataset. Each unit contains the average of the distance to all neighboring units. Metacluster divisions are overlaid. Areas of high distance correspond primarily to a metacluster division. (b) Density map for the ATAC-seq SOM. The color corresponds to the number of chromatin regions found in each unit. Metacluster divisions are overlaid. Most metaclusters are ruled by a few high density units.

**Supplementary Figure 8.**
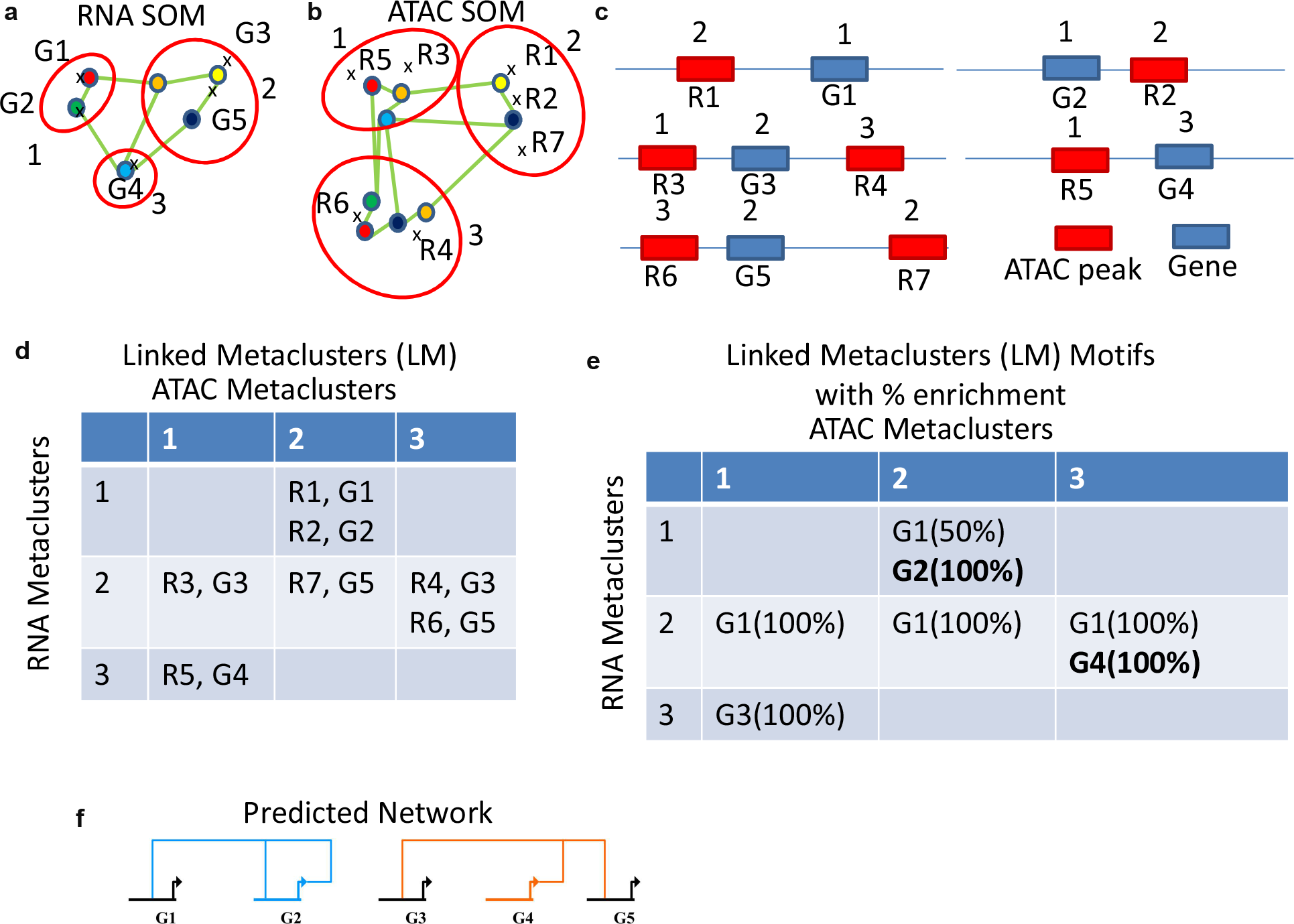
SOM Linking Overview. (a) An example SOM after training on RNA-seq data. Metaclusters 1, 2, and 3 contain genes (G1, G2), (G3, G5), and (G4) respectively. (a) An example SOM after training on ATAC-seq data. Metaclusters 1, 2, and 3 contain genome regions (R3, R5), (R1, R2, R7), and (R4, R6) respectively. (c) An example of how the genes in (a) and the genome regions in (b) could be arranged with their respective metaclusters. (d) The final list of linked metaclusters (LM) that result from the above system. Note that Region 1 and 2 both end up in the same LM (ATAC 2, RNA 1) because they are both in ATAC metacluster 2 and their nearby genes, G1 and G2, are both in RNA metaclusters 1. (e) Example motif enrichments for each gene in (a) in each LM. Bolded genes have a significant enrichment over the background. G1 is found too highly in many LMs and might have an extremely permissive motif. In LM (ATAC 1, RNA 3), G3 motif is found, but would not be called significant due to it being only 1 observation. (f) An example gene regulatory network generated from (e).

**Supplementary Figure 9.**
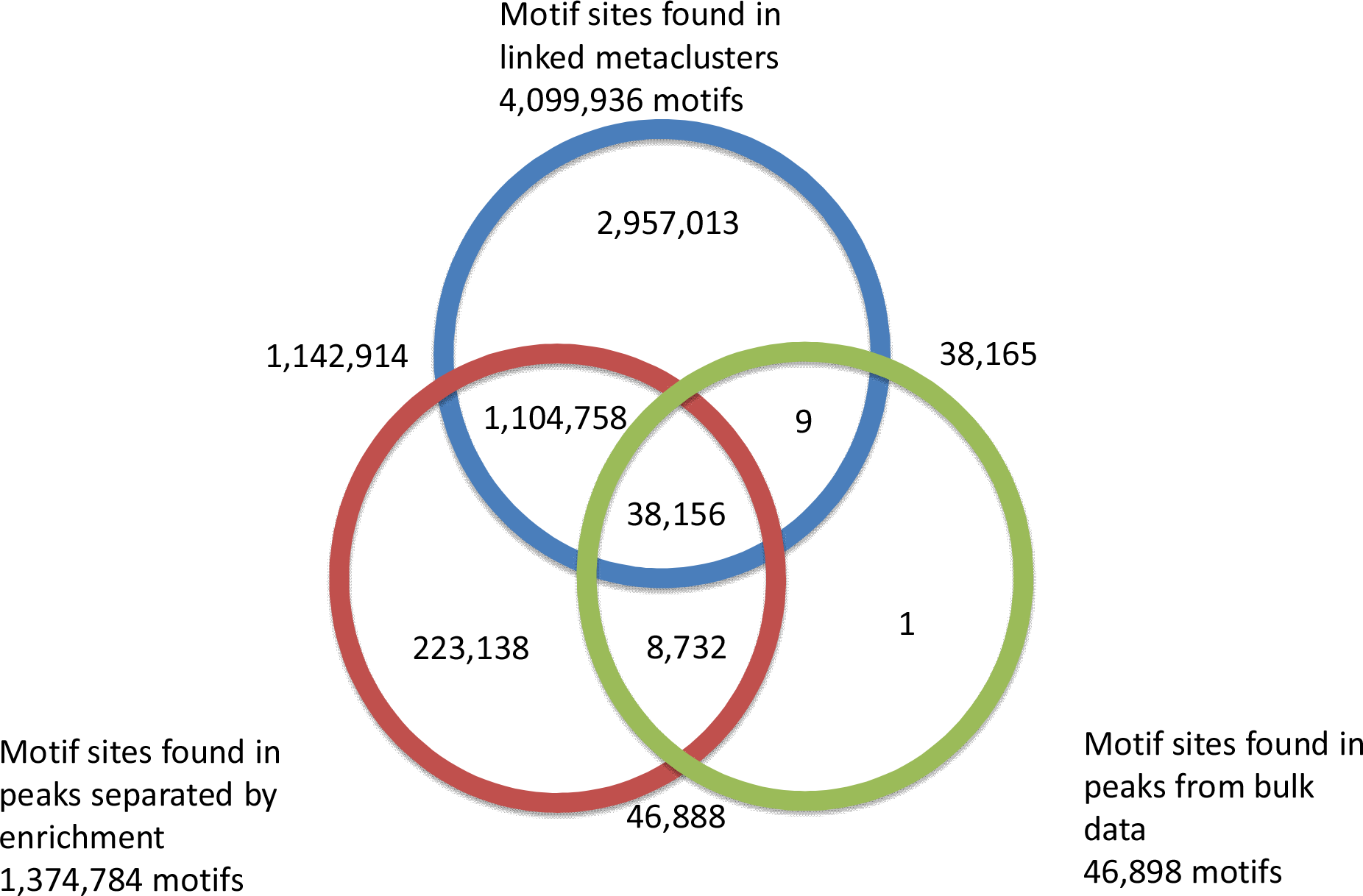
Motif mining efficiency using various techniques. Graph detailing the number of motifs found using the same set of peaks with different groupings using the same q-value<.05 cutoff.

**Supplementary Figure 10.**
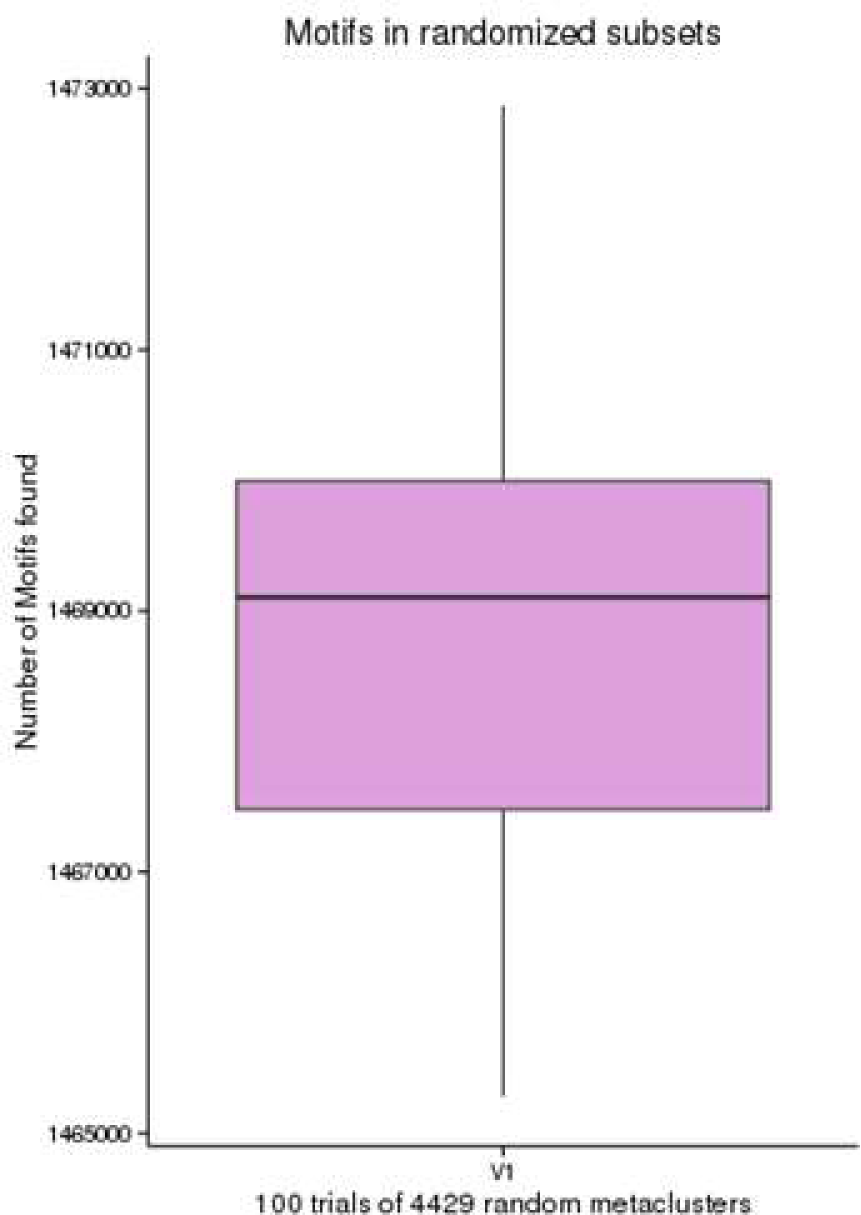
Motif scanning statistics for random separation validation. The distribution of motifs found by randomly splitting the peaks from Supp. Fig. 6 into 103 × 43 = 4,429 synthetic linked metaclusters(LM). The mean was ~1,469,000 motifs which is significantly fewer than the ~4 million found in the real LMs.

**Supplementary Figure 11.**
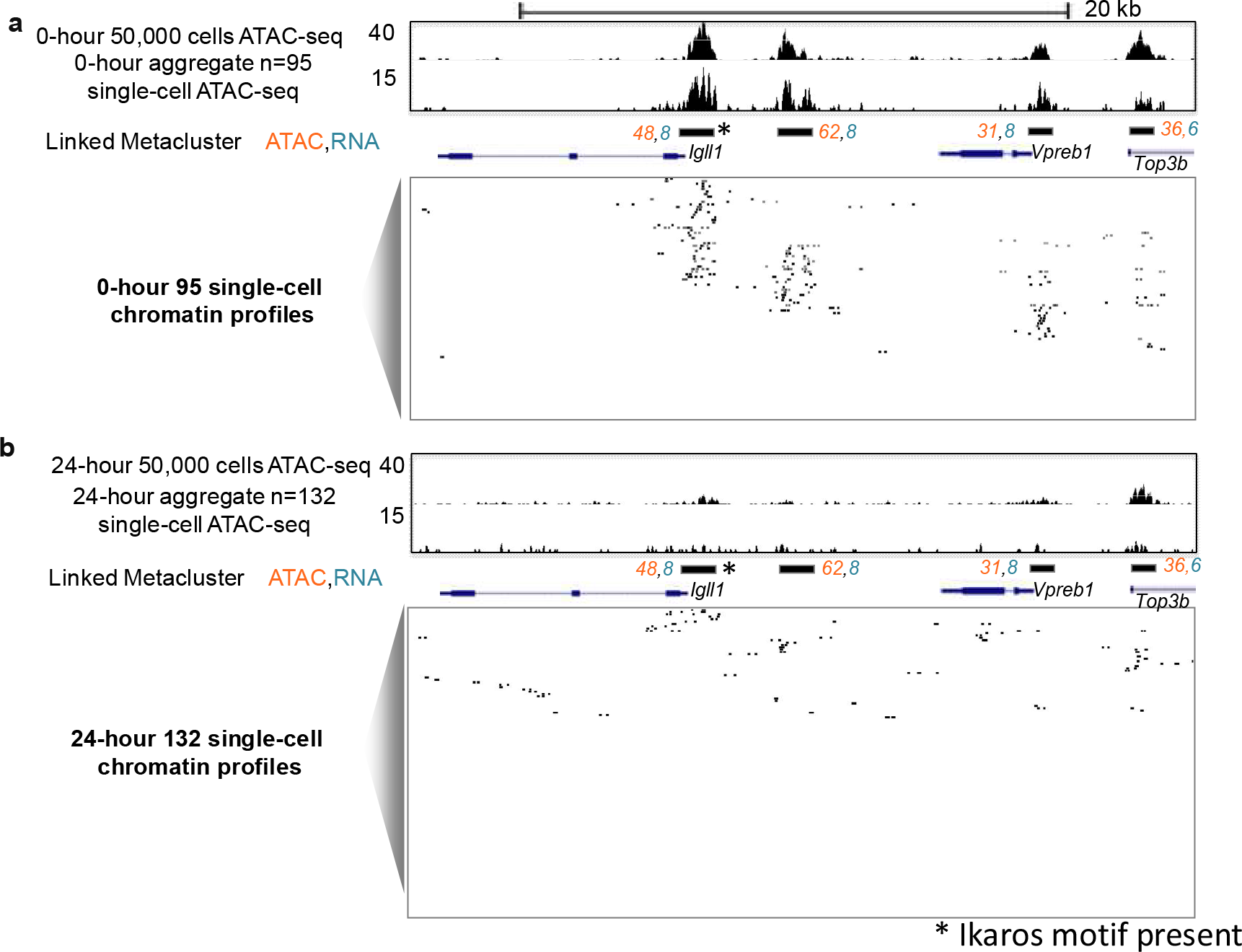
Chromatin accessibility patterns around *Igll1* and *Vpreb1* locus revealed by scATAC-seq labeled by SOMatic. (a-b) UCSC genome browser screenshots of the *Igll1* and *Vpreb1* loci with bulk (50,000 cells), aggregate (95 single-cells averaged) and single-cell ATAC-seq for 0 (a; 95 single-cells) and 24-hour (b;132 single-cells) pre-B cells. Linked SOM ids (ATAC, RNA) are depicted for all chromatin elements.

**Supplementary Figure 12.**
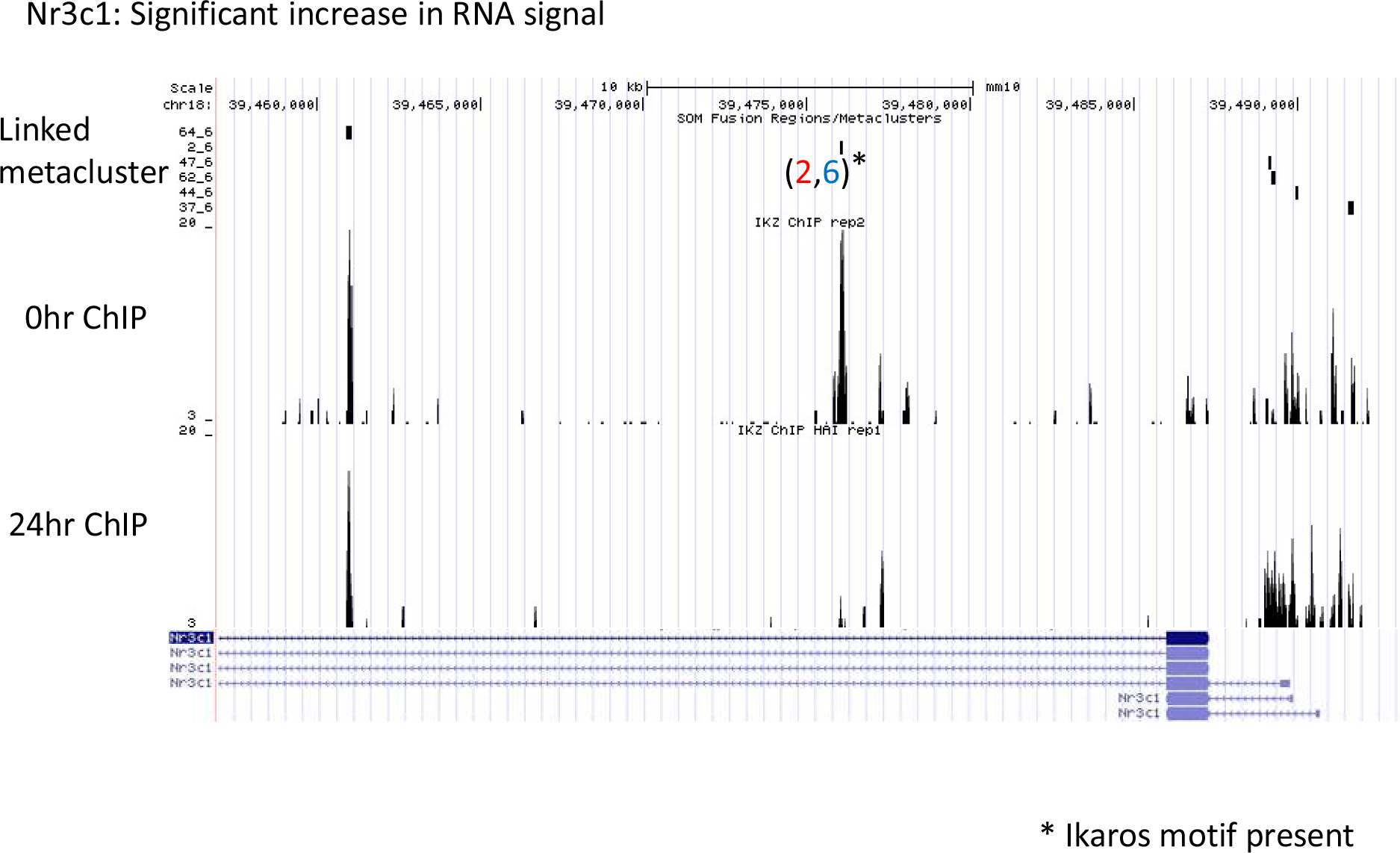
ChIP-seq validation of Ikaraos binding near Nr3c1. UCSC genome browser snapshots of Ikaros ChIP data taken at the 0-hour and 24-hour timepoints near *Nr3c1*. The location of the predicted motif is noted along with its linked metacluster ID. The marked location has a significant change in binding at the marked location over the time course.

**Supplementary Figure 13.**
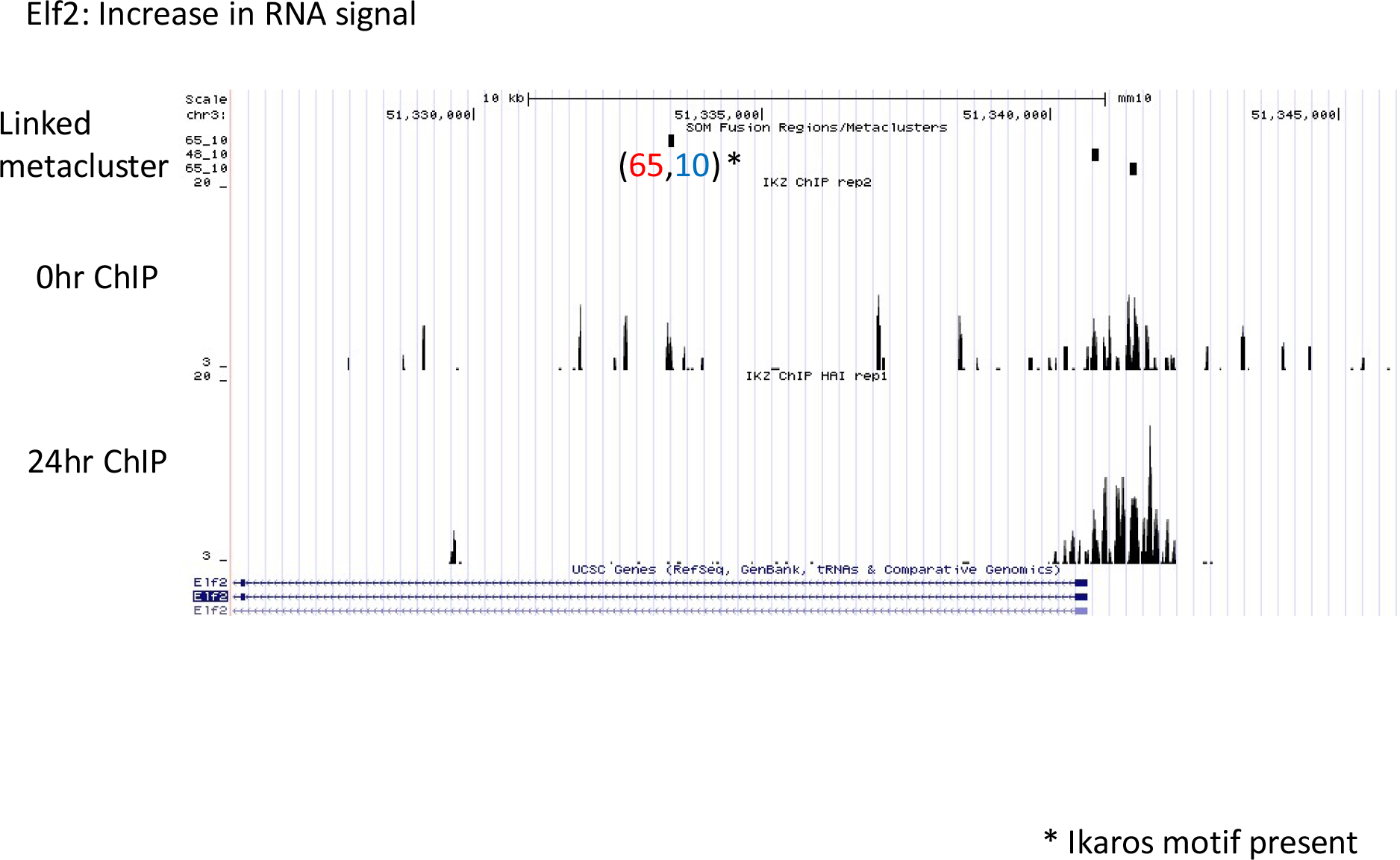
ChIP-seq validation of Ikaraos binding near *Elf2*. UCSC genome browser snapshots of Ikaros ChIP data taken at the 0-hour and 24-hour timepoints near *Elf2*. The location of the predicted motif is noted along with its linked metacluster ID. The marked location has a significant change in binding at the marked location over the time course.

**Supplementary Figure 14.**
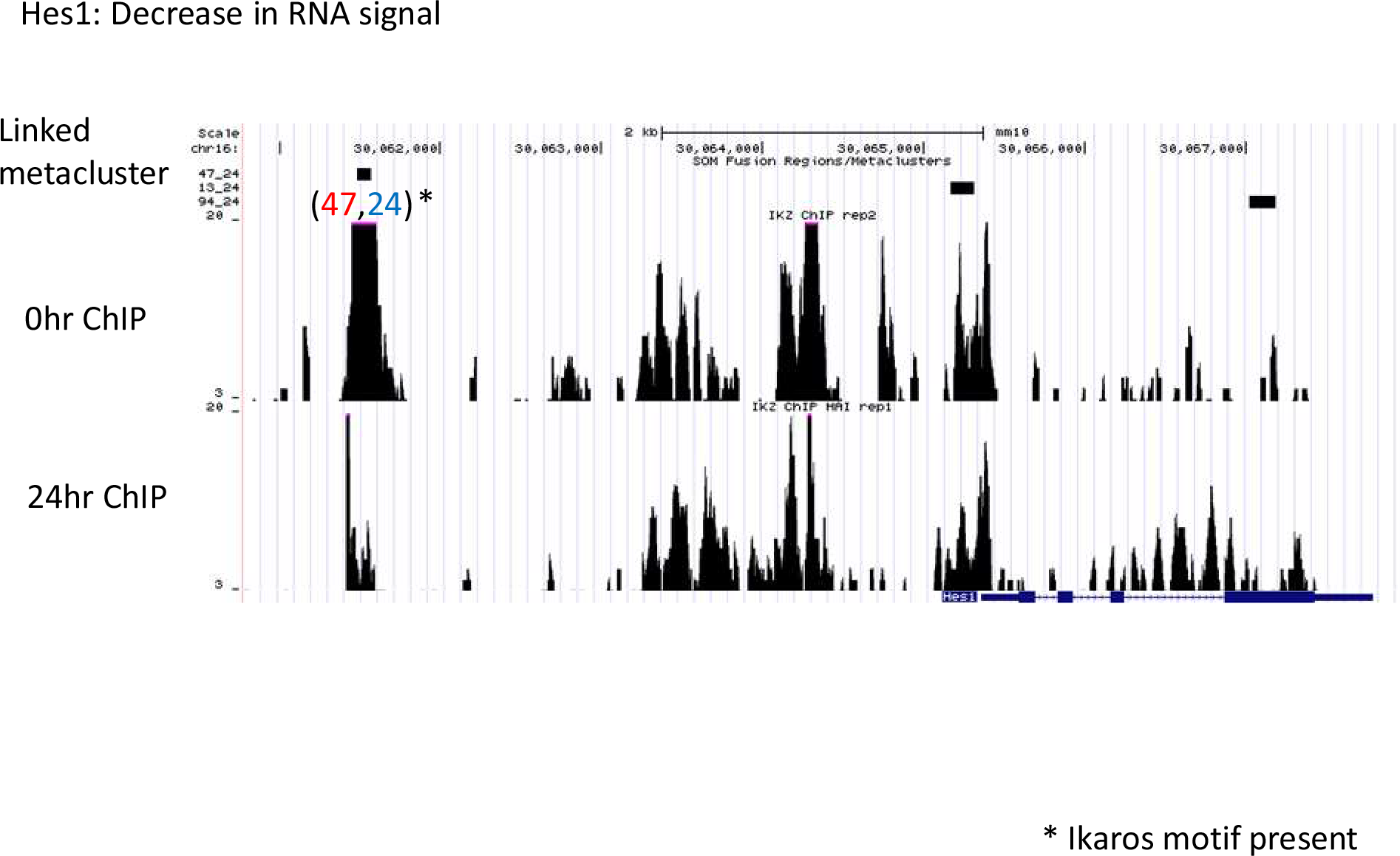
ChIP-seq validation of Ikaraos binding near *Hes1*. UCSC genome browser snapshots of Ikaros ChIP data taken at the 0-hour and 24-hour timepoints near *Hes1*. The location of the predicted motif is noted along with its linked metacluster ID. The marked location has a significant change in binding at the marked location over the time course.

**Supplementary Figure 15.**
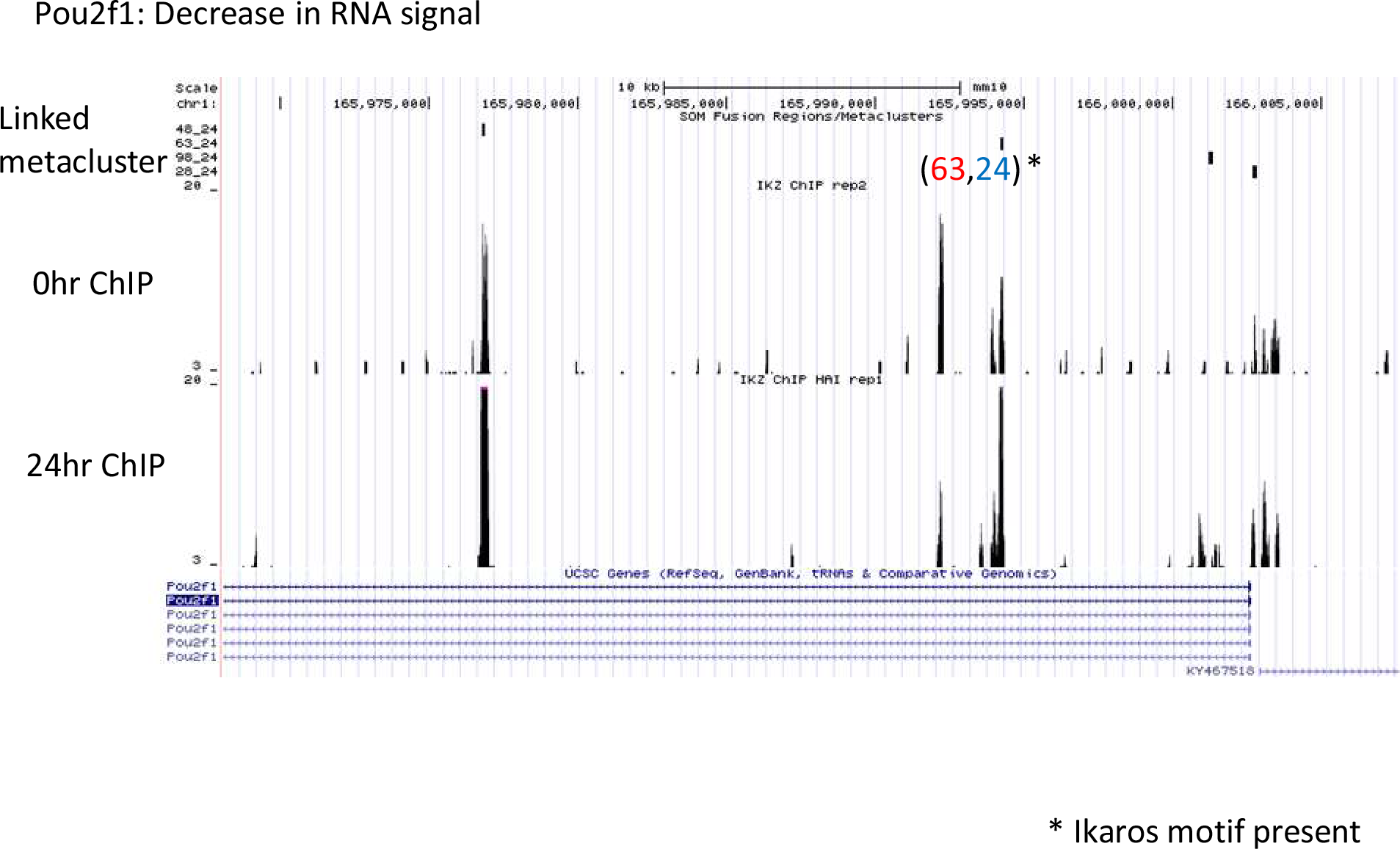
ChIP-seq validation of Ikaraos binding near *Pou2f1*. UCSC genome browser snapshots of Ikaros ChIP data taken at the 0-hour and 24-hour timepoints near *Pou2f1*. The location of the predicted motif is noted along with its linked metacluster ID. The marked location has a significant change in binding at the marked location over the time course.

**Supplementary Figure 16.**
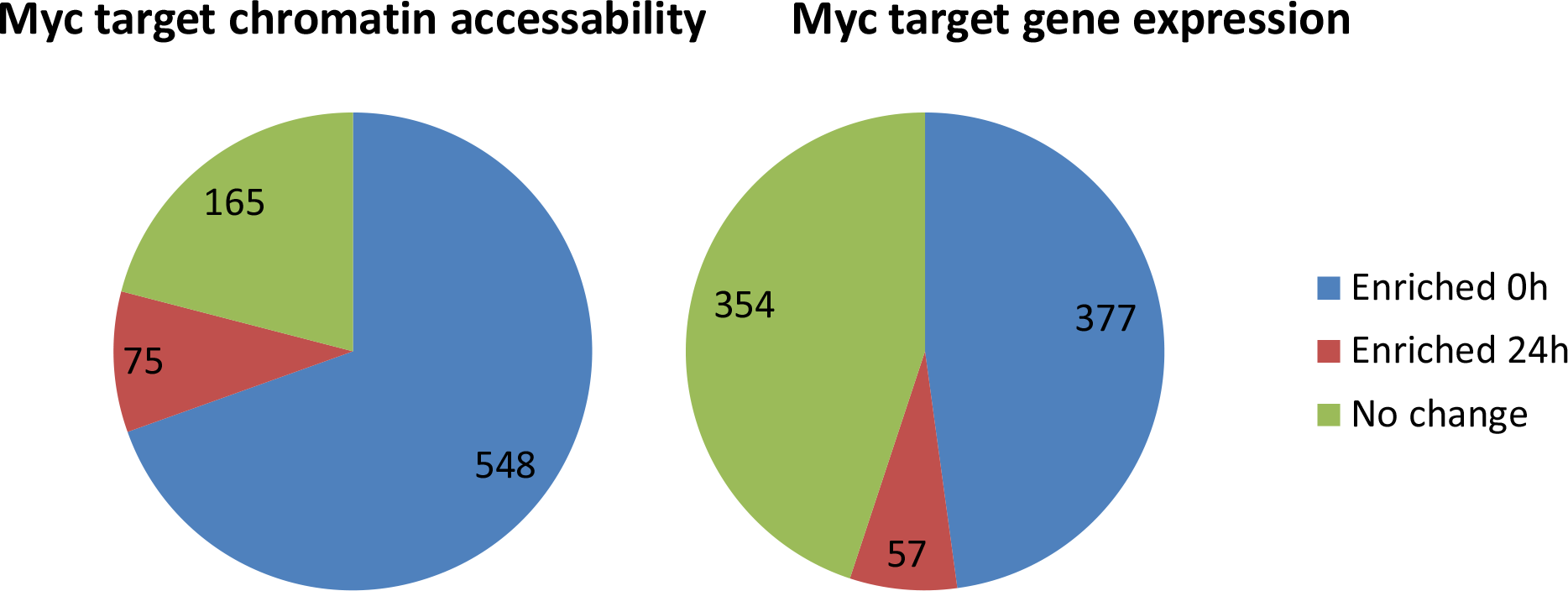
Downstream *Myc* target gene expression and chromatin accessibility dynamics. Myc (whose signal drops dramatically from 0- to 24-hour) downstream targets were predicted in a method similar to that in Figure 4. Around half of these react with a drop in signal with a small portion reacting with an increase. This is similar to the change in chromatin accessibility at the predicted binding sites near these genes.

**Supplementary Figure 17.**
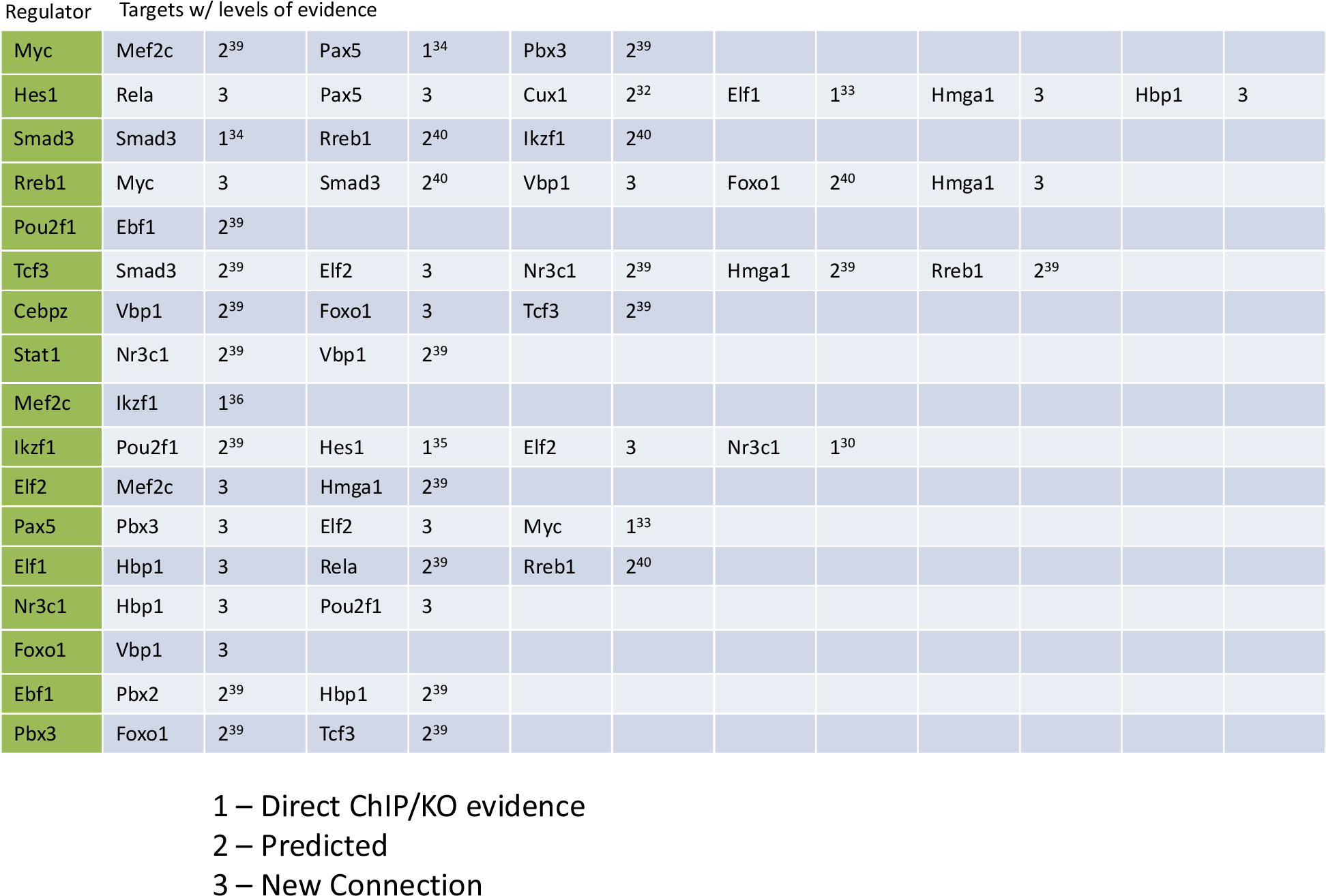
Gene regulatory connections downstream of Ikaros with levels of known evidence. A list of transcription factors with significant changes over the time course and the transcription factors were predicted to regulate. Each regulated gene is followed by a label for the level of existing evidence and reference number if relevant.

